# Neuronal heterogeneity in the medial septum and diagonal band of Broca: classes and continua

**DOI:** 10.1101/2024.08.27.609593

**Authors:** Felix Kuhn, Petra Mocellin, Stefano Pupe, Lihua Wang, Andrew L. Lemire, Liudmila Sosulina, Oliver Barnstedt, Nelson Spruston, Mark S. Cembrowski, Stefan Remy

## Abstract

The medial septum and diagonal band of Broca (MSDB) is known for its diverse populations of cholinergic, GABAergic, and glutamatergic neurons, each contributing to various cognitive processes. However, cell-specific manipulations within this region often result in incongruous behavioral outcomes and reach conflicting conclusions, likely because of a hitherto unknown molecular complexity in its cellular landscape. In this study, we employed single-cell RNA sequencing to thoroughly describe the heterogeneity of MSDB neurons. We confirmed previously established neuronal classes, found gene expression gradients within them and revealed genetically defined subclusters. Moreover, we characterized the genetic profiles of these neuronal subclusters and mapped their spatial distribution. Our analysis presents a comprehensive description of the heterogeneity of MSDB neurons, provides marker genes to target them, explains previous paradoxical results, and opens unexplored avenues to study the impact of neuromodulators in the basal forebrain.

## INTRODUCTION

Located in the medial-ventral portion of the basal forebrain, the medial septum and diagonal band of Broca (MSDB) is a part of the limbic system, highly interconnected with the hippocampal formation and the hypothalamus^1^. Behavioral studies on this brain region trace back to the early 1950s^2,3^, revealing a multifaceted role of the septal area. Its functions include the regulation of food and water intake^4,5^, navigation and memory^6,7^ and the support of reinforcing behaviours^3,8^. Technical progress in identifying, recording, and manipulating specific neuronal types within defined brain regions, combined with advances in behavioral quantification^9,10^, has resulted in an increasingly complex – and often contradictory – picture of MSDB cell types and their role in behavior.

Despite a broad consensus regarding the presence of three primary cell classes^11–13–^ cholinergic, GABAergic, and glutamatergic neurons – molecular, electrophysiological, and functional studies suggest a broader heterogeneity. For example, the MSDB GABAergic neurons were initially thought to be characterized by the expression of the calcium-binding protein parvalbumin (*Pvalb*)^14^. However, further investigation revealed diverse firing patterns among the GABAergic population^15,16^, suggesting the existence of at least two different subpopulations. Additionally, some MSDB neurons that are positive for calretinin (*Calb2*), a marker for GABAergic neurons in the hippocampal formation^17^, do not colocalize with *Pvalb*^18^. The resulting complexity also limits the usefulness of widely used genetic mouse lines targeting GABAergic neurons such as VGAT-Cre, Gad2-Cre, PV-Cre and SST-Cre^19–22^, as each involves only insufficiently characterized subsets of GABAergic neurons.

Similarly, cholinergic and glutamatergic MSDB neurons are likely also more heterogeneous than previously appreciated. A recent study revealed that two subpopulations of cholinergic neurons within the MSDB exist, can be differentiated based on their expression of calbindin (*Calb1*), and play distinct roles in behavior^23^. Additionally, manipulation of the MSDB glutamatergic population targeting specific projections resulted in diverse behavioral outcomes, including aversion^24^, feeding^25^, reinforcement and exploration^26,27^. Previous work in subiculum^28^, ventromedial prefrontal cortex^29^, and primary visual cortex^30^ suggests that projection target heterogeneity is often accompanied by molecular heterogeneity, further hinting at molecular diversity within the MSDB populations.

By employing single-cell RNA sequencing (scRNA-Seq) and spatial transcriptomics, we aimed at describing the molecular profile and spatial distribution of the MSDB neurons. Our analyses not only confirm the existence of three major classes of cholinergic, GABAergic, and glutamatergic neurons, but also identify previously undescribed subclusters and novel markers to interrogate them.

## RESULTS

### Transcriptomic profiling of the mouse medial septal area

To study the diverse neuronal population of the MSDB we performed single cell RNA sequencing (scRNA-Seq). The region of interest was microdissected based on anatomical landmarks and the cells were manually dissociated (n=4 mice, 2 batches). After the exclusion of non-neuronal cell types and low-quality cells, our dataset comprised a total of 903 MSDB neurons (median number of features per cell: 5563 – interquartile range (IQR): 4477-6947; median RNA count per cell: 154711 - IQR: 89072-273657; median percentage of mitochondrial genes: 2.7% - IQR: 2.26-3.35 %, Figures S1A-C)

To identify the main neuronal classes, we clustered the data using the Leiden algorithm^31^. The most informative resolution was determined using a co-dependency index (CDI)-based differential expression analysis^32^ (Figures S1D-F). Four transcriptionally distinct classes emerged from the clustering which constitute four separate groups on a Unifold Manifold Approximation and Projection (UMAP) on a two-dimensional plane (Figure 1A). Since the MSDB is populated by cholinergic, glutamatergic, and GABAergic neurons^12,13,33^, we examined the expression of commonly used genetic markers for each of these cell classes (Figure 1B). We identified one cholinergic class characterized by the expression of the neurotransmitter-synthesizing enzyme choline acetyltransferase (*Chat*) as well as the choline transporter *Slc5a7*; one glutamatergic class selectively expressing the gene *Slc17a6* for the vesicular glutamate transporter type 2 (VGluT2); and two GABAergic classes with high expression levels of the gene *Slc32a1* coding for the vesicular GABA transporter (VGAT). Despite the broad use of glutamate decarboxylase 1 (*Gad1*) and glutamate decarboxylase 2 (*Gad2*) Cre mouse lines to target MSDB GABAergic neurons^19,34–36^, we found that both markers were not specific. Instead, *Gad1* was broadly expressed also in the cholinergic class, while *Gad2* expression spanned across all neurons.

**Figure 1.**
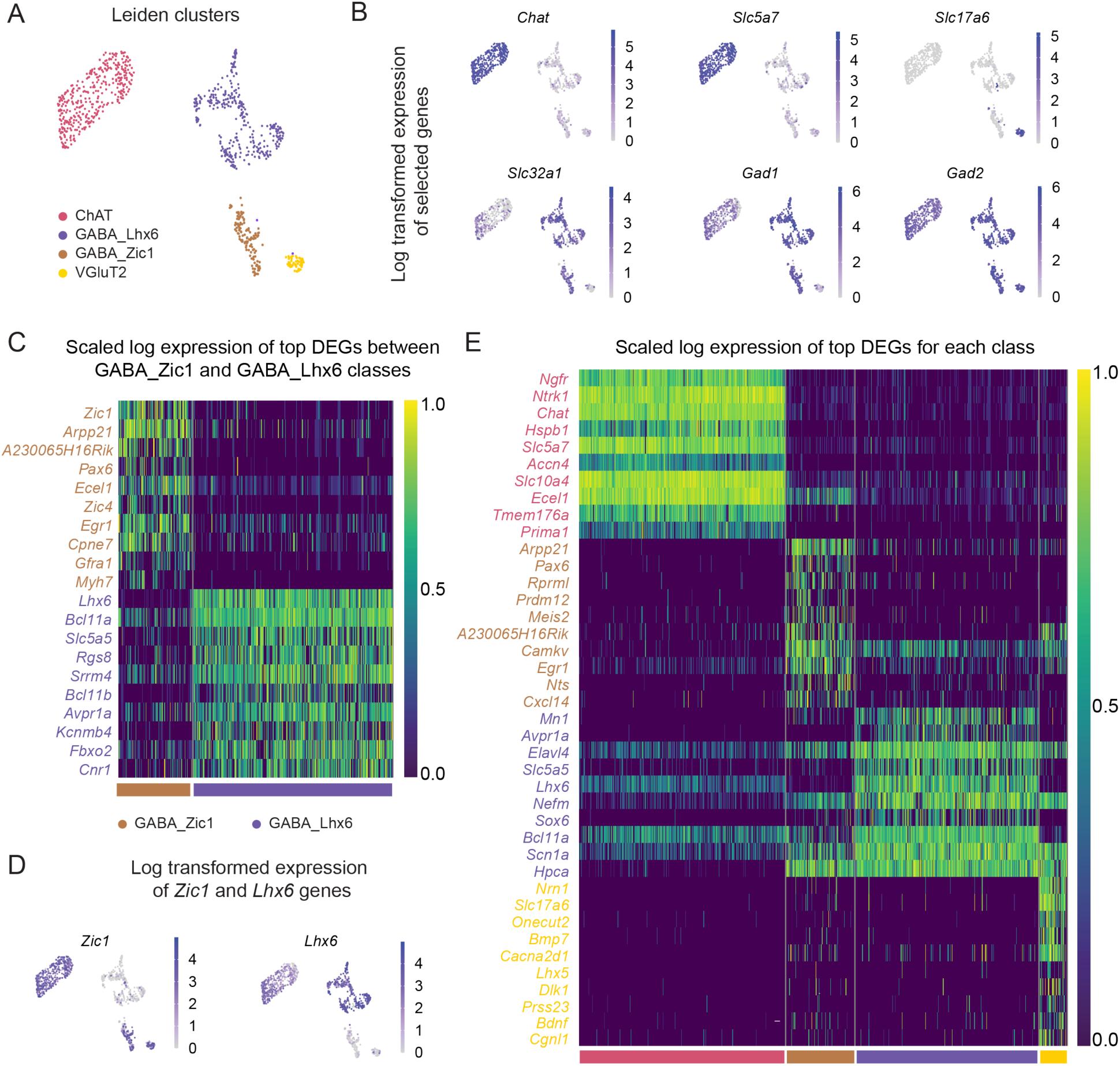
scRNA-Seq analysis of the mouse septal area. A) Leiden clusters with resolution of 0.1. B) Feature plots showing the log-transformed expression of classic cholinergic (*Chat, Slc5a7*), GABAergic (*Slc32a1, Gad1, Gad2*), and glutamatergic (*Slc17a6*) markers in the MSDB. C) Heat map representing the scaled log-transformed expression of DEGs between GABA_Zic1 and GABA_Lhx6 clusters sorted by false discovery rate (FDR-) corrected p-value (Wilcoxon rank sum test). D) Feature plots of log-transformed expression of *Zic1* and *Lhx6* genes. E) Heat map representing the scaled log-transformed expression of the top 10 DEGs for each cluster sorted by FDR-corrected p-value (Wilcoxon rank sum test). Only genes with a log2 fold-change expression (log2FC) higher than 1.5 and that had a difference in the fraction of expression of at least 30% in the respective cell class compared to the rest of the neuronal population were considered as DEGs in Figure 1C and 1E. Classes color code throughout the figure: ChAT in pink, GABA_Lhx6 in purple, GABA_Zic1 in brown, VGluT2 in yellow.

To differentiate between the two GABA classes, we conducted a differential expression analysis using Wilcoxon rank sum test. The analysis revealed that *Zic1* and *Lhx6*, two transcription factors involved in the septal region development^37,38^, are mutually exclusive in the two GABAergic classes (Figures 1C-D). Building upon this analysis, we named the four neuron classes ChAT, VGluT2, GABA_Lhx6 and GABA_Zic1, respectively. To provide a detailed overview of gene expression differences across all neuron classes, we identified the ten most significant differentially expressed genes (DEGs) for each of them in comparison with all MSDB neurons outside of the respective clusters using Wilcoxon rank sum test (Figure 1E).

### MSDB scRNA-Seq integration with spatial transcriptomic datasets

While scRNA-Seq is the state-of-the-art method for identifying and classifying neuronal subpopulations within a specific region, it cannot capture the cells’ spatial distribution and often exhibits bias in the relative proportions of different cell types^39^. To quantify the abundance of the neuronal populations and investigate their localization in the MSDB, we combined our scRNA-Seq analysis with the analysis of publicly available spatial transcriptomic datasets obtained using Multiplexed Error-Robust Fluorescence *In Situ* Hybridization (MERFISH). Specifically, we used the Zhuang-ABCA-1 and Zhuang-ABCA-2^40^ datasets, which comprise mouse brain coronal sections targeting 1,122 genes, available from the Allen Brain Cell Atlas (ABCA) (Figure S2A). For each coronal slice we analyzed neurons from a selection window in one hemisphere encompassing MSDB^+^ and nearby MSDB^-^ neurons. To identify MSDB^+^ neurons, we refined the segmentation provided with the Zhuang datasets in Allen Mouse Brain Common Coordinate Framework version 3 (CCF) coordinates^41^ with a custom-written tool (see Methods). Our further analysis focused only on neurons, excluding glial, immune, and vascular cells.

Using Canonical Correlation Analysis (CCA)^42^, we predicted the identity of MERFISH neurons in the region of interest using the scRNA-Seq dataset as a reference. To quantify the likelihood of a MERFISH neuron to belong to a scRNA-Seq class, CCA calculated a prediction score ranging from 0 to 1. MERFISH neurons with a prediction score above 0.9 were assigned to their predicted class. We further compared marker gene expression profiles of individual classes between scRNA-Seq and MERFISH datasets, ensuring consistency (Figure S2B).

ScRNA-Seq analysis may not capture all the cell types in a given brain region, due to biological and technical limitations^43^. To confirm that the scRNA-Seq data included all MSDB neuronal classes present in the MERFISH dataset, we projected the cells of both MERFISH and scRNA-Seq datasets onto a UMAP in a joint feature space (jFS) (Figure S2C) and quantified the number of neurons for each class in both datasets (Figures 2A - scRNA-Seq, 2B - MERFISH). We found that 90% of the MERFISH neurons are assigned to one of the four scRNA-Seq classes with a prediction score above 0.9. Thus, there is a high correspondence between the two datasets with regard to MSDB^+^ neuronal classes. Moreover, the 10% MERFISH neurons with a prediction score lower than 0.9 did not segregate into a separate group in the UMAP visualization. This indicates that our scRNA-Seq analysis reliably captured all main cell classes in the MSDB, suggesting that the lower prediction scores were likely due to imperfect single-cell segmentation or cell overlap in the MERFISH data rather than the presence of unrecognized neuronal classes.

**Figure 2.**
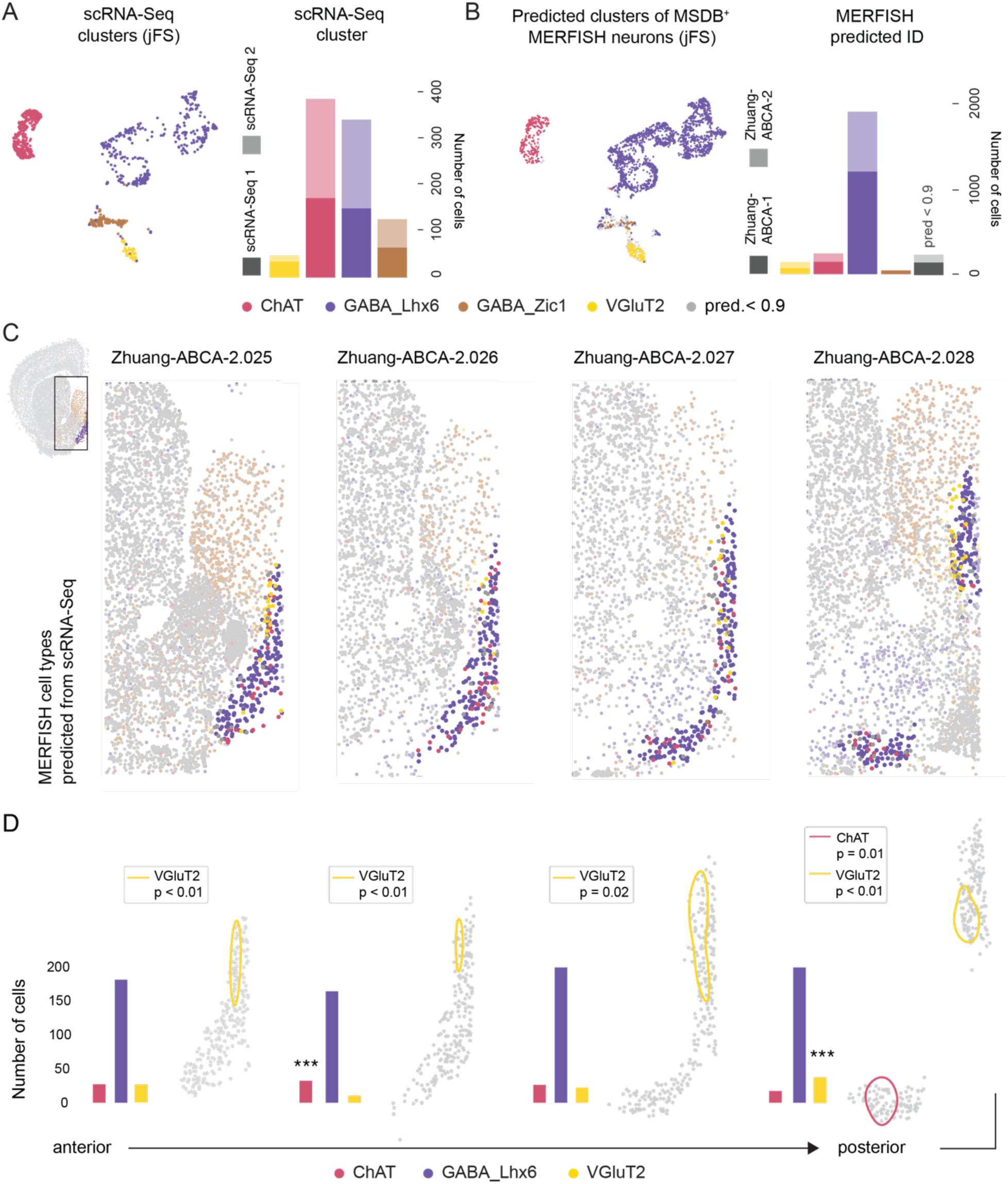
Integration of the scRNA-Seq and the MERFISH ABCA datasets reveals the spatial distribution of four neuronal clusters. A) UMAP in jFS with scRNA-Seq neurons color-coded by cluster identity (left panel) and the proportion of neurons assigned to each cluster from each experimental batch (sc-RNA-Seq1 and scRNA-Seq2, respectively; right panel). B) UMAP in jFS with MERFISH neurons color-coded by predicted identity (left panel) and the proportion of neurons assigned to each cluster including neurons with a prediction score < 0.9 (right panel). C) Coronal slices of the Zhuang-ABCA-2 MERFISH dataset (slices 025, 026, 027, and 028) with cells color-coded based on the predicted identity from the scRNA-Seq dataset (MSDB^+^ neurons in bold colors, MSDB^-^ neurons in faint colors). Scale bars correspond to 5 CCF units for both axes. D) Full width at half maximum (FWHM) of the maximum-likelihood kernel density estimate (ml-KDE) for VGluT2 and ChAT cells when they show significant spatial clustering inside the MSDB in the respective slice (for significance calculation, see Methods). Bar plots represent the number of neurons per cluster for each coronal slice. Stars represent significant enrichment of the respective cluster on the respective slice (* p<0.05, ** p<0.01, *** p< 0.001). Scale bars correspond to 5 CCF units for both axes. Classes color code throughout the figure: ChAT in pink, GABA_Lhx6 in purple, GABA_Zic1 in brown, VGluT2 in yellow, MERFISH neurons with prediction scores < 0.9 in gray.

Next, we mapped all the neurons with a prediction score above 0.9 in the spatial domain and marked each cell with the corresponding class-based color. For the visualization of the cell position, we used four representative slices along the anterior-posterior extent of the MSDB in both Zhuang datasets (Figures 2C, S3A). Consistent with previous studies, the GABA_Lhx6 population was localized in the MSDB^33^, while GABA_Zic1 neurons were mostly distributed in the lateral septum (LS)^44,45^. We concluded that the scRNA-Seq *Zic1^+^* neurons in our scRNA-Seq dataset belong to the transition area between MSDB and LS, given the lack of distinct anatomical demarcations between the two^46^. Here, we focus our in-depth analysis of MSDB GABAergic neurons on the GABA_Lhx6 class only.

Leveraging the spatial resolution provided by the MERFISH datasets, we employed a Monte-Carlo-based shuffling approach to detect significant spatial clustering of the MSDB classes along the anterior-posterior axis. The analysis revealed that GABA neurons are homogeneously distributed, ChAT neurons are more prominent in the anterior sections, and VGluT2 neurons populate more abundantly the posterior part (Figures 2D, S3B). Furthermore, we examined whether there was significant spatial clustering along the dorsoventral and mediolateral axes by determining significant spatial trends in individual coronal sections using a Monte-Carlo based permutation test of global Moran’s I^47^ (Figures 2D, S3B, see Methods). We found that the abundance of VGluT2 neurons is significantly enriched in the dorso-lateral regions of the MSDB and that in posterior regions of the MSDB the abundance of ChAT neurons is significantly enriched in the diagonal band of Broca.

### Cholinergic neurons in the MSDB are organized in a transcriptomically continuous cluster with two distinct endpoints

For many decades MSDB cholinergic neurons have been studied as the main source of acetylcholine to the hippocampus^12^, involved in memory^48^, attention^49^, and type II theta oscillations^50^. However, further studies showed cholinergic subpopulations co-expressing other neurotransmitters^51^, projection targets outside the hippocampal formation^52^, and multiplex behavioral outcomes^23^, suggesting a broader heterogeneity than previously acknowledged.

Using Leiden clustering, we aimed to explore whether MSDB cholinergic neurons form molecularly distinct subclusters. A CDI-based criterion determined an optimal clustering resolution of 0.4, resulting in two subclusters (Figures 3A, S4A-C). To explore the transcriptional features distinguishing the two subclusters, we conducted differential gene expression analysis (Figure 3B). *Gal* and *Amigo2* appeared among the most significant DEGs between the two ChAT clusters, thus we named the clusters ChAT_Gal and ChAT_Amigo2, respectively.

**Figure 3.**
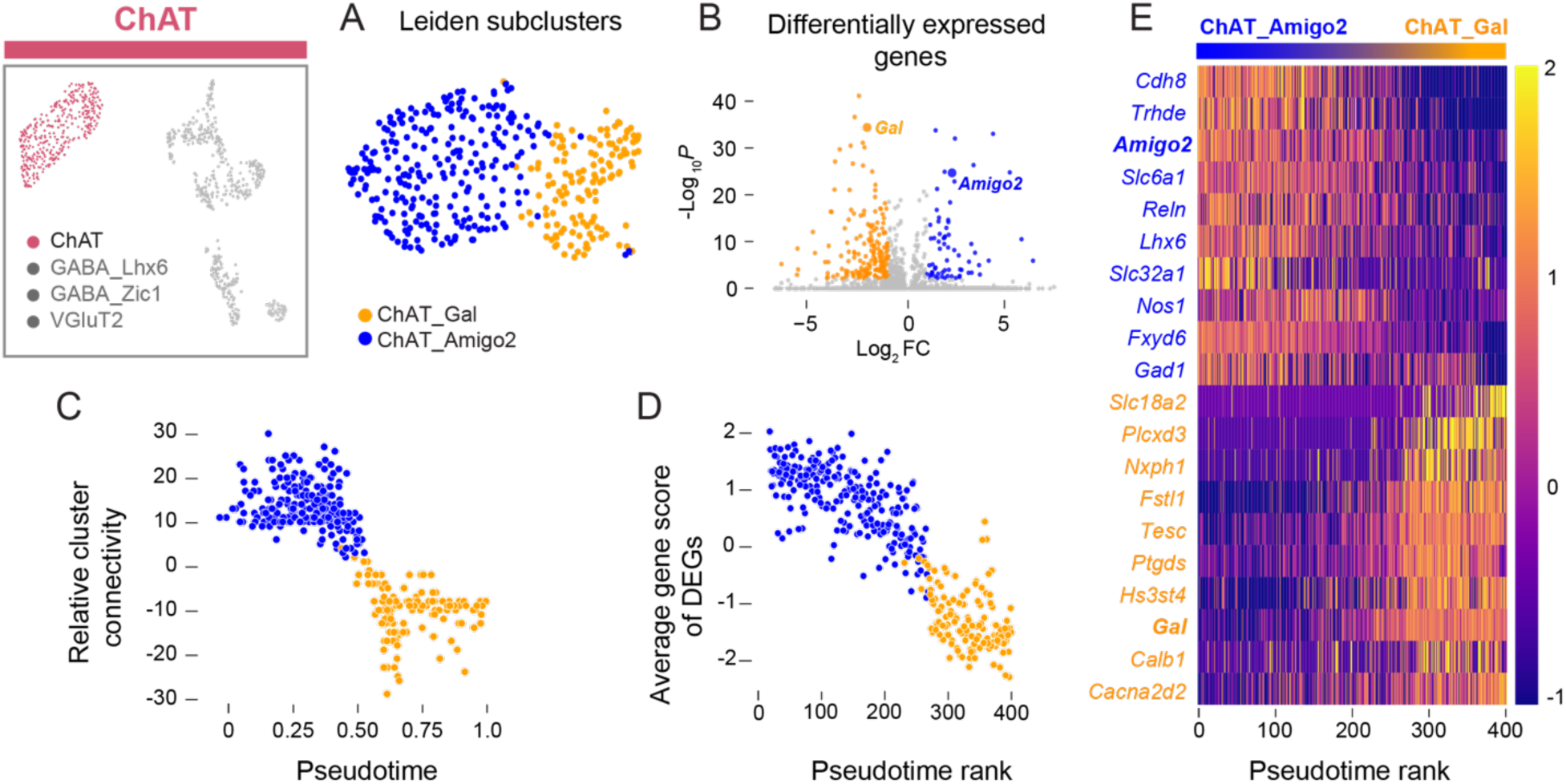
Characterization of the MSDB cholinergic cluster. A) UMAP of cholinergic subclusters (ChAT_Gal in orange and ChAT_Amigo2 in blue) from Leiden sub-clustering with resolution 0.4. B) Volcano plot of differentially expressed genes between the two subclusters. Gray dots have either an absolute log2FC value below 1 or an FDR-corrected p-value below 0.001 (bold dots indicate *Gal* (orange) and *Amigo2* (blue)). C) Relative cluster connectivity of neurons as a function of diffusion pseudotime. D) Cumulative differential expression score per cell of the 20 most significant DEGs per cluster (FDR corrected p-value was below 10^-5^ for all genes) ordered by diffusion pseudotime rank. E) Heatmap showing the standardized log-transformed expression values of selected DEGs in MSDB^+^ cholinergic cells. Cells are ordered by diffusion pseudotime rank and genes by absolute log2FC value.

To investigate if these two clusters represent individual classes or are part of a transcriptomic continuum, we utilized the gap c statistic^53^. This relies on two metrics: (1) the diffusion pseudotime metric^54^ and (2) the relative cluster connectivity metric. Diffusion pseudotime measures transitions between cells using diffusion-like random walks such that cells with a similar transcriptomic profile lie closer on a pseudotime axis compared to cells with more distinct profiles. The relative cluster connectivity describes how confidently a cell can be assigned to one of two different clusters. We calculated diffusion pseudotime for the MSDB ChAT neurons along the direction of highest variance between the two ChAT subclusters, as well as their relative cluster connectivities. Our analysis neither revealed significant gaps along the diffusion pseudotime axis, nor the relative cluster connectivity axis (Figure 3C). Additionally, we examined cumulative differential expression scores^53^ of the 20 most significant DEGs between the two clusters, sorted according to their rank on the pseudotime axis. Notably, we also did not observe any significant gap in the cumulative expression scores (Figure 3D). Taken together, these findings suggest that ChAT neurons in the MSDB are organized in a transcriptomic continuum with two distinct endpoints.

To further characterize the ChAT continuum, we plotted the expression profiles of a selection of significant DEGs for each endpoint along the diffusion pseudotime axis (Figure 3E). We discovered that ChAT_Amigo2 neurons highly express genes related to the synthesis and transport of the neurotransmitter GABA (*Slc6a1*, *Gad1*, *Slc32a1*), while ChAT_Gal neurons are more closely linked to neuromodulators as demonstrated by the presence of the neuropeptide galanin (*Gal*) and the vesicular monoamine transporter 2 (*Slc18a2* – VMAT2) that facilitate dopamine and serotonin loading into synaptic vesicles. These results align with recent evidence of a ChAT-GABA co-releasing population in the MSDB^51,55,56^. In line with the notion of a continuum, the expression profile of the DEGs does not show a clear gap between the two ChAT subclusters, as expression ranges of individual genes vary along the diffusion pseudotime axis.

Next, we studied the spatial distribution of the MSDB^+^ ChAT neurons and investigated whether they are genetically distinct from cholinergic neurons populating the nearby Striatum (ChAT::Striatum). First, we visualized the scRNA-Seq and the MERFISH datasets of the ChAT MSDB^+^ neurons on a UMAP in jFS and predicted the subcluster identity of the MERFISH cells using CCA (Figures 4A, S4D). We then repeated this integration, this time also including cells from inside a selection window around the MSDB including MSDB^+^ neurons, as well as neurons from the nearby Striatum. We found that ChAT::Striatum neurons are transcriptomically distinct from MSDB^+^ ChAT neurons (Figure 4B). Furthermore, we found that the cholinergic cells of the scRNA-Seq dataset mainly colocalize with the MSDB^+^ neuron cluster on the UMAP in jFS, highlighting the proper alignment between ChAT neurons of our scRNA-Seq dataset and MSDB^+^ neurons in the MERFISH dataset. In line with previous literature, differential expression analysis revealed that ChAT::Striatum neurons are significantly enriched in genes for the dopamine receptor *Drd2*^57^, the homeodomain transcription factor *Gbx2*^58^, and the protein phosphatase *Ppp1r1b*^59^ (Figure 4C).

**Figure 4.**
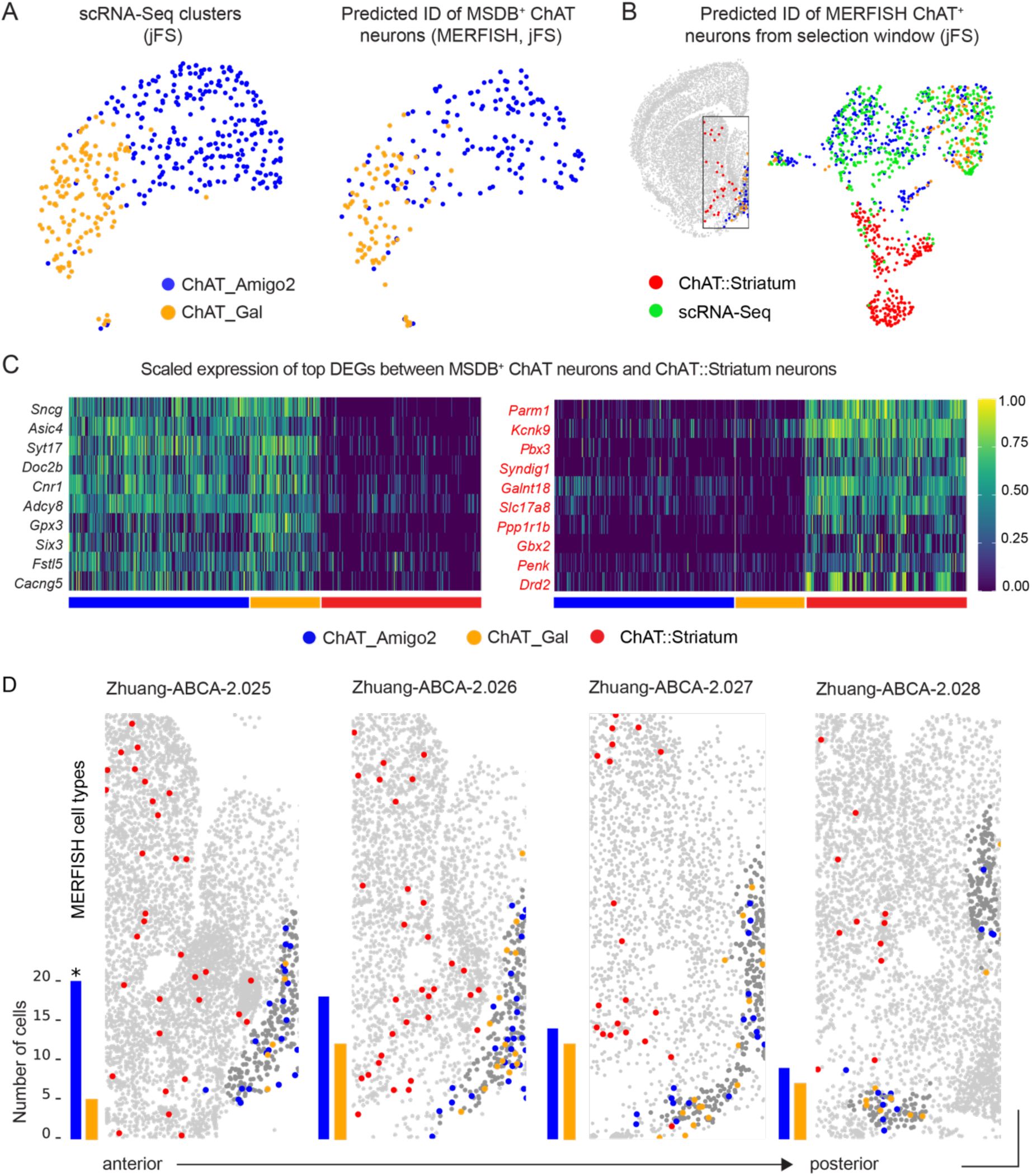
Spatial distribution of cholinergic MSDB neurons and their relation to striatal ChAT neurons. A) UMAPs in jFS of MSDB^+^ ChAT neurons from the scRNA-Seq and MERFISH datasets (left panel: scRNA-Seq, right panel: MERFISH); color code represents Leiden subclusters (scRNA-Seq) and predicted cluster identities (MERFISH). B) UMAP of ChAT neurons from MSDB scRNA-Seq dataset and ChAT MERFISH neurons from the selection window in jFS. C) Heat map of the scaled log-transformed expression of selected DEGs of ChAT MSDB^+^ neurons and ChAT::Striatum neurons. D) Spatial distribution of the ChAT clusters along the anterior-posterior axis. (ChAT_Amigo2 in blue, ChAT_Gal in orange, ChAT::Striatum in red). Bar plots represent the abundance of the two MSDB^+^ ChAT neuronal populations in each slice. The star represents significant enrichment of the ChAT_Amigo2 subcluster in the most anterior slice following Monte-Carlo based permutation testing (p_value = 0.04). Scale bars correspond to 5 CCF units for both axes.

Additionally, we investigated the spatially distribution of the MSDB^+^ ChAT_Gal and ChAT_Amigo2 neurons, as well as the ChAT::Striatum neurons in the selection window for both MERFISH datasets. Monte-Carlo based spatial enrichment analysis among MSDB ChAT neurons uncovered significant enrichment of ChAT_Amigo2 neurons in anterior regions of the MSDB (Figures 4D, S4E).

### Heterogeneity of the MSDB Lhx6 GABAergic class

GABAergic neurons in the MSDB have been extensively studied due to their dense projections to the hippocampal formation^14^. In particular, it was found that MSDB *Pvalb* interneurons expressing the hyperpolarization-activated, cyclic nucleotide-gated non-selective cation (HCN) channel exert pacemaking activity, capable of synchronizing both the local network and the hippocampal formation during theta oscillations^16,60–62^. However, these neurons represent only a fraction of the GABAergic cell population in the MSDB, and there is no clear consensus on the overall number of distinct GABAergic clusters and their characteristics.

To address this issue, we sub-clustered the GABA_Lhx6 class. Using Leiden clustering with a resolution of 1.2 as determined by the CDI-based method, we delineated six distinct subclusters (Figures 5A, S5A-D). To investigate whether these subclusters are transcriptionally distinct or part of a continuum, we used the gap c statistics and found that each GABA_Lhx6 subcluster blends into multiple other GABA_Lhx6 subclusters continuously (Figures 5B, S5E).

**Figure 5.**
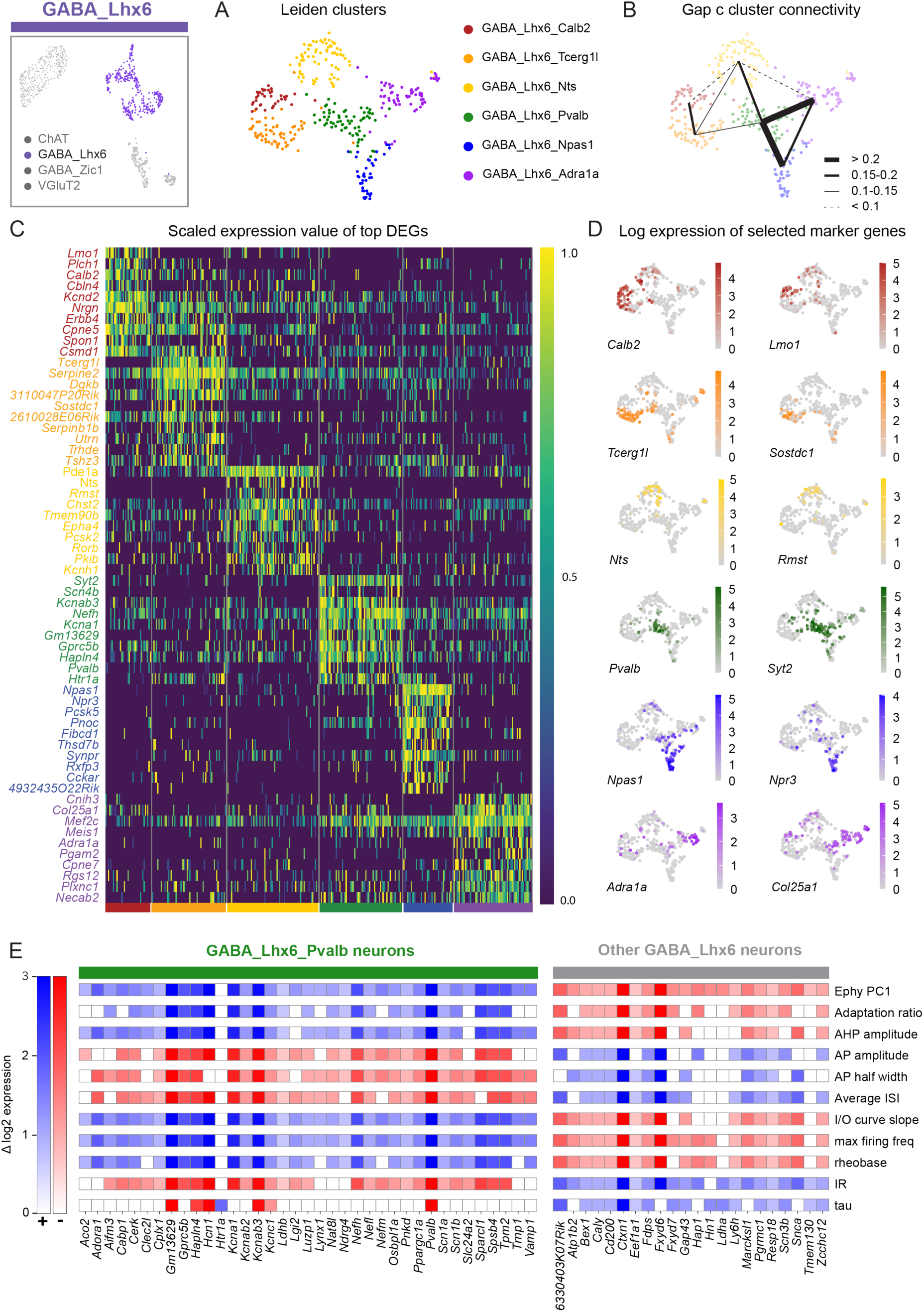
Transcriptomic profile of the GABA_Lhx6 class. A) UMAP of GABAergic Lhx6 subclusters from Leiden sub-clustering with resolution 1.2. B) Strength of cluster connectivity based on gap c statistic (the higher the value, the smoother the transition from one cluster to the other). C) Heatmap showing the scaled log-transformed expression value of top DEGs per Leiden subcluster ranked by FDR-corrected p-value (Wilcoxon rank sum test). Only genes with log2FC value higher than 1.5 and that had a fraction of expression which was 30 percent higher in the subcluster of interest compared to the rest of the population were considered as DEGs D) Feature plots showing the log-transformed expression of selected marker genes per Leiden subcluster. E) Log-transformed differential expression of positive (green) and negative (gray) marker genes of the GABA_Lhx6_Pvalb subcluster which are positively (blue) or negatively (red) correlated with identified electrophysiological properties based on the Bomkamp et al. database^63^.

To characterize genetic profiles for the GABAergic subclusters, we identified the 10 most significant DEGs of each subcluster using the Wilcoxon rank sum test (Figure 5C). We further identified marker genes with high specificity based on CDI, including well-known genes associated with MSDB cell types or common Cre-lines (Figure 5D). Based on the expression profiles of these genes, we named the subclusters as follows: GABA_Lhx6_Calb2, distinguished by high expression of the gene encoding the calcium-binding protein calretinin (*Calb2*); GABA_Lhx6_Tcerg1l, marked by elevated expression of the gene encoding the transcription elongation regulator 1 (*Tcerg1l*); GABA_Lhx6_Nts, characterized by the presence of the gene for the endogenous neuropeptide neurotensin (*Nts*); GABA_Lhx6_Pvalb, identified by the expression of the gene encoding the calcium-binding protein parvalbumin (*Pvalb*); GABA_Lhx6_Npas1, marked by the expression of the gene for the Neuronal PAS Domain Protein 1 (*Npas1*); and GABA_Lhx6_Adra1a, characterized by the expression of the gene encoding the adrenergic receptor alpha 1A (*Adra1a*).

Since MSDB *Pvalb* neurons play a key role in hippocampal theta rhythm, and given the extensive *in vivo* and *in vitro* recordings studying their features^16,60^, we explored potential correlations between gene expression and electrophysiological properties in GABA_Lhx6_Pvalb^+^ and GABA_Lhx6_Pvalb^-^ neurons. For this reason, we compared differentially expressed genes in the GABA_Lhx6_Pvalb subcluster with electrophysiological properties significantly linked to these genes. Our analysis is based on a publicly available database by Bomkamp and colleagues^63^ correlating gene expression gradients of inhibitory neurons with gradients in electrophysiological properties (Figure 5E). We found that the GABA_Lhx6_Pvalb^+^ subcluster is enriched in genes associated with high firing frequency, high rheobase, low input resistance, and narrow action potential width (Figure 5E). Indeed, these electrophysiological features are classically associated with MSDB *Pvalb* neuron activity^64^, further validating our findings.

Next, we integrated MERFISH and scRNA-Seq neurons of the GABA_Lhx6 class using CCA-based integration. By predicting the subcluster identity of GABAergic MERFISH neurons and mapping the prediction scores onto a UMAP in jFS (Figure 6A), we observed high prediction scores, except for neurons situated between two clusters. This occurrence reflects the concept of continuous clusters: neurons positioned between two endpoints of a continuum are equally assigned to both clusters, resulting in a prediction score of 0.5. For further analysis of the GABA_Lhx6 class, we included all MERFISH neurons with maximum prediction scores equal or above 0.5. To confirm the alignment between scRNA-Seq clusters and MERFISH GABAergic clusters we also projected MSDB^+^ scRNA-Seq neurons from the GABA_Lhx6 class onto the UMAP in jFS (Figure 6B). We additionally subpartitioned the MERFISH neurons only using Leiden Clustering and projected the clusters in the jFS UMAP (Figure 6C). We found each MERFISH subcluster region on the jFS UMAPs to consist mainly of MERFISH neurons belonging to a specific subcluster as predicted with the scRNA-Seq dataset. This indicates proper alignment between the transcriptomic profile of GABAergic MERFISH and scRNA-Seq neurons. Additionally, we confirmed that selected marker genes for GABAergic subclusters in the scRNA-Seq data exhibit consistent expression patterns with the corresponding subclusters in the MERFISH data (Figure S6A).

**Figure 6.**
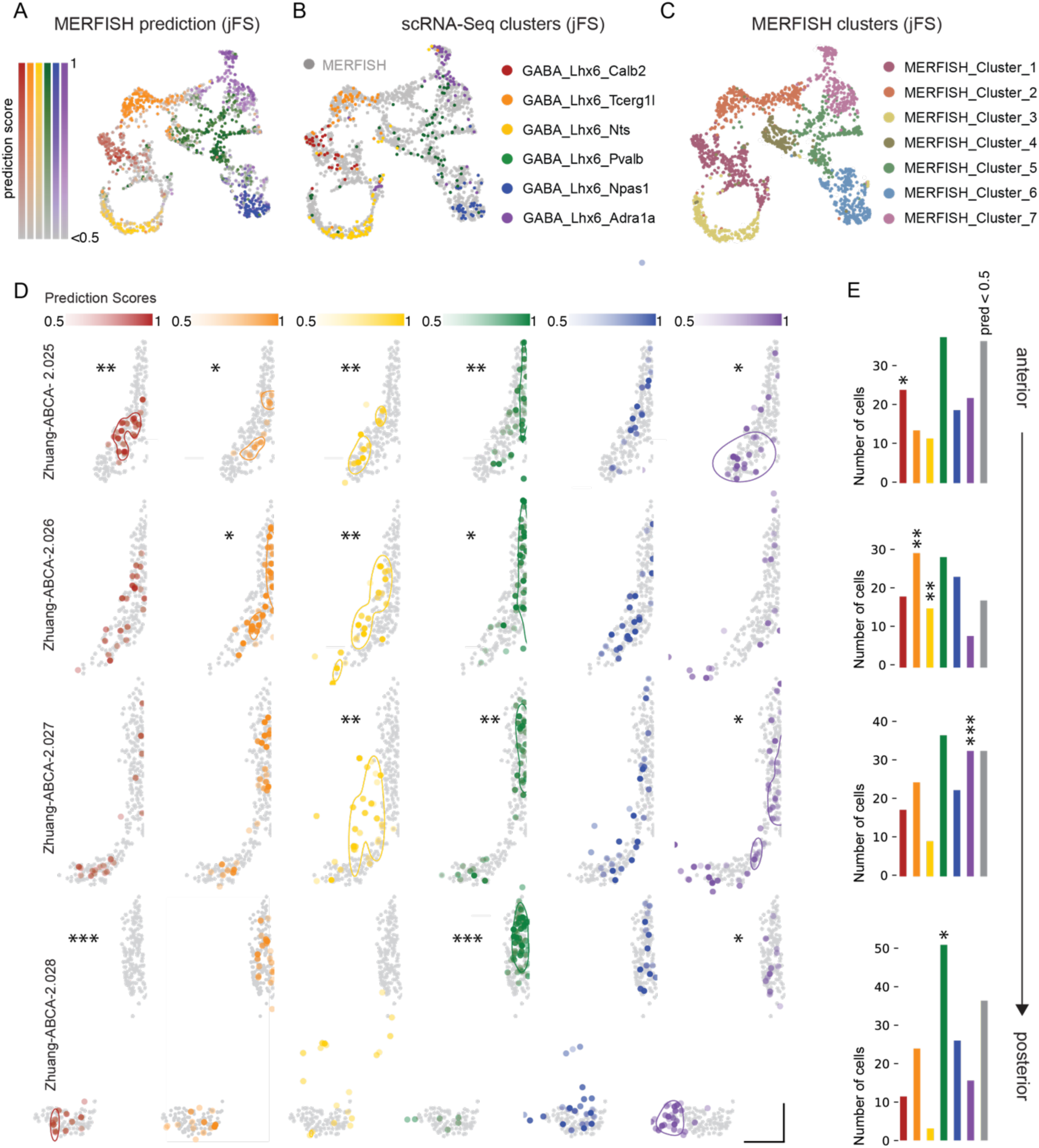
Spatial distribution of GABA_Lhx6 subclusters. A) UMAP in jFS of scRNA-Seq and MERFISH GABA_Lhx6 neurons with MERFISH neurons color-coded by prediction identity and prediction score. B) UMAP in jFS of the GABA_Lhx6 cluster with scRNA-Seq neurons color-coded by subcluster identity and MERFISH neurons in gray. C) UMAP in jFS of MERFISH neurons color coded by MERFISH subcluster identity (Leiden resolution = 0.3). D) Spatial expression patterns of predicted MERFISH subclusters color-coded by prediction score in the Zhuang-ABCA-2 dataset. If significant spatial patterns were found inside of MSDB by the Monte-Carlo approach (* p<0.05, ** p<0.01, *** p< 0.001; see Methods) the FWHM of the respective ml-KDE was plotted. Scale bars correspond to 5 CCF units for both axes. E) Number of neurons per cluster for each coronal slice. Stars represent significant enrichment of the respective cluster on the respective slice as calculated by the Monte-Carlo approach (* p<0.05, ** p<0.01, *** p< 0.001).

In order to determine spatial trends within each coronal slice and along the anterior-posterior axis of the MSDB, we mirrored the Monte-Carlo based analysis previously employed for the main MSDB neuronal classes and for the ChAT neurons, this time shuffling across GABAergic neurons only. Our analysis unveiled that GABA_Lhx6_Pvalb neurons predominantly occupy the midline of the MSDB, and GABA_Lhx6_Nts neurons being the most lateral and extending into the surrounding areas. In the most posterior slice, GABA_Lhx6_Calb2 and GABA_Lhx6_Adra1a preferentially occupy the diagonal band of Broca with GABA_Lhx6_Adra1a also extending further into the surroundings of the diagonal Band (Figures 6D, S6B). Additionally, we observed that GABA_Lhx6_Nts, GABA_Lhx6_Tcerg1l, and GABA_Calb2 are significantly enriched in the anterior to middle portions of the MSDB, while GABA_Lhx6_Adra1a and GABA_Lhx6_Pvalb are significantly enriched in middle to posterior regions (Figures 6E, S6C).

### MSDB^+^ VGluT2 neurons are transcriptomically distinct

Accounting for a quarter of MSDB neurons, VGluT2 cells have been found to form the smallest neuronal subset in this brain region^11^. Our scRNA-Seq data reflects this, capturing only a limited number of VGluT2^+^ neurons in the MSDB. Although the VGluT2 class in the scRNA-Seq dataset appeared to be homogeneous, its size was insufficient for a representative sub-clustering analysis, like the one conducted for the GABA and ChAT classes. Nonetheless, we aimed to further characterize this cell class given the recent interest in its functional role^13,26,27,65,66^. Therefore, we investigated how the MSDB VGluT2 neurons identified in our scRNA-Seq dataset differ from other VGluT2 cells in the septal surroundings. To achieve this, we used Leiden clustering on the different MERFISH VGluT2 neurons within a defined spatial selection window. Our analysis revealed four main VGluT2 clusters (Figure 7A).

**Figure 7.**
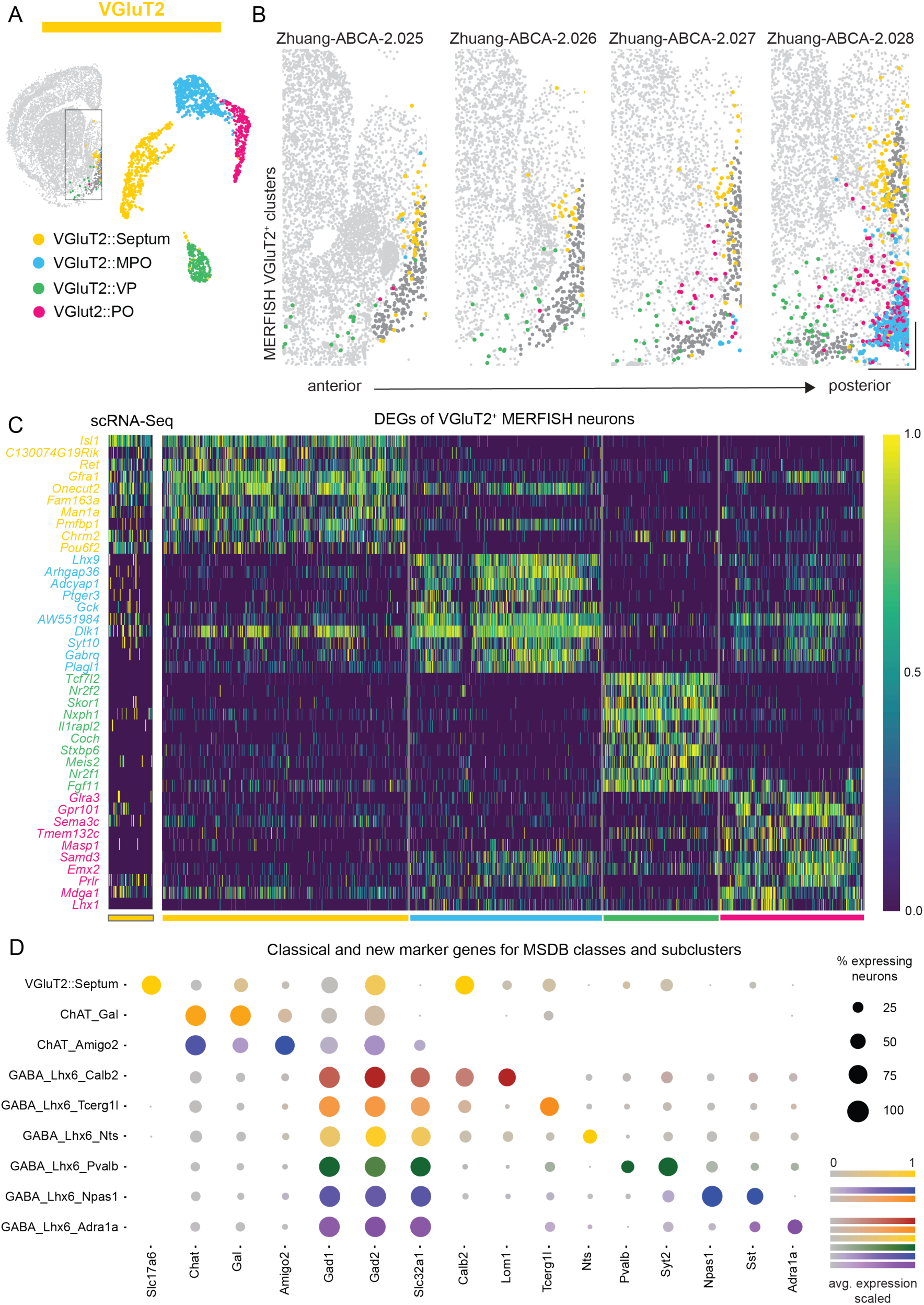
Transcriptomic profile of MSDB^+^ and MSDB^-^ VGluT2 neurons and expression profiles of selected genes across all MSDB subclusters. A) Selection window and UMAP displaying the clusters of VGluT2^+^ neurons from a spatial selection window in the two MERFISH datasets. B) Spatial distribution of VGluT2 neurons in the selection window along the anterior-posterior axis of the Zhuang-ABCA-2 dataset. Scale bars correspond to 5 CCF units for both axes. C) Heatmap of the scaled log-transformed expression of DEGs for each MERFISH VGluT2 cluster and corresponding expression in the scRNA-Seq VGluT2 neurons. D) Expression profile of selected scRNA-Seq marker genes for each class and subcluster. The dot size represents the percentage of cells expressing a given gene for each subcluster. The color scale represents the average expression of the respective gene, scaled to the highest value across subclusters.

When observing the spatial distribution of the VGluT2 clusters (Figures 7B, S7A), only one of them was overlapping with the septal area, thus we named it VGluT2::Septum. A group of VGluT2^+^ neurons was located in the ventral and lateral portions of the MSDB, overlapping with the medial and lateral preoptic area (VGluT2::PO). Another population appeared very dense in the ventro-medial portion of the more posterior slices, specifically overlapping with the medial preoptic area (VGluT2::MPO). Finally, a fourth group of VGluT2 neurons was located laterally, and was mainly co-localized with the ventral pallidum (VGluT2::VP). Next, we investigated the DEGs for each of these MERFISH VGluT2 clusters and compared the expression profiles of their most significant DEGs with their expression profiles in scRNA-Seq VGluT2^+^ neurons (Figure 7C). This approach confirmed that the VGluT2 neurons in our scRNA-Seq dataset share their transcriptomic profile with the neurons of the VGluT2::Septum MERFISH cluster, expressing the transcription factor *Onecut2* and the proto-oncogene *Ret*^67,68^, among the others. In line with previous studies, we detected high levels of the transcription regulators *Tcf7l2* and *Skor1* in the VGluT2::VP neurons^69,70^, *Lhx1* in the VGluT2::PO neurons^71^, as well as specific expression of the prolactin receptor (*Prlr*) in the VGluT2::MPO neurons^72^. Notably, VGluT2::Septum neurons are situated dorsally within the septum, occupying an area that, based on the expression of the GABA_Lhx6 and GABA_Zic1 clusters, overlaps with both the medial and lateral septum. Moreover, VGluT2 neuron clusters increasingly intermingle with one another along the anterior-posterior axis, with VGluT2::VP cells almost completely overlapping with the diagonal band of Broca in the more posterior parts of the MSDB (Figures 7B, S7A).

Finally, we present a dot plot (Figure 7D) showcasing the average expression of both classical and newly identified marker genes. Our analysis reveals that newly identified markers are not only subcluster specific within their class, but also across MSDB classes. The sole exception is *Calb2*, which exhibits a high expression profile also in the glutamatergic class. Instead, *Lmo1* emerges as a valuable marker to target the GABA_Lhx6_Calb2 subcluster specifically. The dot plot also emphasizes the non-specific expression of the classical GABAergic markers *Gad1* and *Gad2*. This underscores the significance of this study in identifying more precise promoters to target the MSDB populations. Here, we provide novel insights into the transcriptomic landscape of the MSDB, offering a foundation for future research aimed at selectively targeting subcluster-specific populations in functional experiments.

## DISCUSSION

The cellular landscape of the MSDB has been extensively studied, yet a consensus on the number and characteristics of its neuronal populations has remained elusive. Combining scRNA-Seq and MERFISH datasets, our work provides a comprehensive analysis of the transcriptomic profile and spatial distribution of the MSDB neurons. First, we confirmed the presence of distinct classes for cholinergic, GABAergic, and glutamatergic neurons. Then, we sub-clustered them and identified their genetic expression profiles. We analyzed the spatial distribution of neuronal classes and their subclusters within each coronal slice along the MSDB anterior-posterior axis. Additionally, we compared transcriptomic differences between MSDB^+^ and nearby MSDB^-^ neurons expressing the same neurotransmitter.

We found that the MSDB cholinergic class is organized along a gene expression continuum with two distinct endpoints: one exhibiting co-expression of genes involved in GABA transport and synthesis (ChAT_Amigo2), and the other associated with neuromodulators such as galanin (ChAT_Gal). Spatial analysis revealed that cholinergic neurons show the highest density in the anterior-middle sections of the MSDB.

The GABAergic population is best described by six subclusters, which blend into each other, highlighting a complex and overlapping landscape. We showed that the genetic profile of the GABA_Lhx6_Pvalb cluster is consistent with the well-documented rhythmically firing *Pvalb*^+^ neurons^16,60^. GABAergic subclusters display specific localization patterns in the MSDB, with GABA_Lhx6_Pvalb neurons concentrated along the midline and GABA_Lhx6_Nts neurons being the most lateralized.

Furthermore, we found the septal VGlut2 neurons (VGluT2::Septum) to be transcriptomically distinct from other basal forebrain (VGluT2::VP) and hypothalamic (VGluT2::MPO, VGluT2::PO) glutamatergic populations. VGluT2::Septum neurons are predominantly localized in the dorso-lateral medial septum and increase in density along the anterior-posterior axis.

### Transcription factor expression in the adult MSDB

The development of the septum follows what has been described as an “onion skin-like” structure^73^. This laminated structure is the result of different streams of neurons that during embryonic days E10.5 to E14.5 migrate to their final location occupying first the medial and then the lateral portion of the septal area^45,73^.

Transcription factors play a crucial role in regulating the neurons’ migration into their final location^74^. Having access to the full transcriptomic profile of the adult septal neurons gave us valuable information about the identity and origin of the different populations. In particular, we could distinguish between lateral and medial septum GABAergic neurons based on the exclusive expression of *Lhx6* and *Zic1*, with the GABA_Lhx6 class populating the MSDB and the GABA_Zic1 class the LS. These findings align with embryonic studies showing the specificity of *Lhx6* and *Sox6* for MSDB neurons, and *Zic* family genes together with *Pax6* for LS GABAergic neurons^33,37^. Our data further confirmed *Zic1* expression in the MS ChAT population^38^, suggesting that MSDB ChAT neurons and LS GABAergic neurons may derive from a common progenitor or are migrating at similar time points. Indeed, recent data highlighted that nearly all cholinergic neurons in the basal forebrain express the vesicular GABA transporter VGAT (*Slc32a1)* at some point in development^51^. As a strong negative marker for MSDB cholinergic neurons, we identified the transcription factor *Gbx2,* which is exclusively expressed in the ChAT::Striatum^58^ population and completely absent in the MSDB ChAT neurons.

Recent studies have highlighted that the *Pvalb^+^* MSDB neurons are generated at E.10.5, being the earliest in the development of MSDB neurons, and thus occupying the most medial portion of the septum^33^. Indeed, our spatial analysis revealed a significant increase of GABA_Lhx6_Pvalb cells around the midline of the MSDB. Our spatial analysis also revealed that the boundary between the MSDB and LS significantly depends on the markers chosen to define it. This suggests that studies using *Pvalb* to identify the MSDB may have excluded more lateral parts, particularly areas populated by VGluT2 neurons.

### Gene expression continuum of septal cholinergic neurons

Being one of the most prominent cholinergic areas in the basal forebrain^75^, and given its direct inputs to the hippocampal formation^76^, multiple lines of research have focused on the properties of MSDB cholinergic neurons. So far, no categorization of cholinergic neuron subtypes in the MSDB exists. While our data confirm that classical markers like *Chat* and *Slc5a7* reliably target the whole MSDB cholinergic population, we demonstrated that cholinergic neurons span a continuum of gene expression.

On one end of the continuum resides the ChAT_Gal population, characterized by high expression levels of galanin. Previous studies described a subpopulation of septal cholinergic neurons expressing this neuropeptide^52,77^ and projecting to the hippocampal formation^78^. Interestingly, it was reported that galanin injection in the MSDB reduces theta amplitude, impairs working memory in the rat dorsal hippocampus^79^, and increases acetylcholine release in the ventral hippocampus^80^. In the hippocampus, it was suggested that galanin had a permissive effect on acetylcholine release, but an excessive release driven by galanin impaired working memory performance^80^. This observation raised some interest for its possible implications in Alzheimer disease (AD): patients were shown to have high levels of galanin in the basal forebrain^81,82^ but it remains unclear to which extent galanin has a neuroprotective or toxic effect on the pathological septo-hippocampal system^83–85^. This might be further complicated by the fact that galanin is expressed, albeit at lower levels, in the glutamatergic and GABAergic neurons of the MSDB. Currently no fluorescence-based reporters are available to study galanin release. Given its potential implication in AD and a so far underexplored role in the regulation of septo-hippocampal activity during learning and memory processes, the development of galanin sensors could strongly advance our understanding of neuropeptide release by the Chat_Gal population. This, in turn, would shed light on both the physiological and pathological effects of galanin on the activity of the MSDB and the hippocampal formation.

On the other end of the gene expression continuum within the cholinergic neuron class, the ChAT_Amigo2 subcluster shows high co-expression of genes involved in GABA synthesis and transport such as *Gad1* (Gad67), *Gad2* (Gad65), *Slc32a1* (VGAT) and *Slc6a1* (GAT1). While we observed that *Gad2* is present in all the MSDB neuronal types, the expression of transporters that are usually located in the presynaptic terminals of GABAergic neurons (GAT1)^86^, or in the GABA containing vesicles (VGAT)^87^ suggests that this population is capable of co-releasing GABA and acetylcholine, as previously reported in functional^55,56,88^ and developmental^51^ studies. Notably, we found that the ChAT_Amigo2 neurons are expressing high levels of reelin (*Reln*), a glycoprotein studied for its role in neurodevelopment^89^, synaptic plasticity^90^, learning^91,92^, and AD^93^. While previous studies have shown reelin positive neurons in the MSDB during development^94^ and in adult mice^95^; and despite the broad interest for reelin and the MSDB in relation to AD^96–98^, no studies to our knowledge have reported its expression in cholinergic neurons or have investigated the functional role of reelin in the MSDB.

### The diversity of MSDB GABAergic neurons

The complexity of the GABAergic cell types in both cortical and subcortical circuits has been widely recognized^99–102^. The GABAergic MSDB neurons were initially studied in relation to the hippocampal formation and *Pvalb* was long considered a key marker for septo-hippocampal GABAergic projections^14^. Subsequently, other GABAergic populations were described based on the observation that some MSDB neurons express GABAergic markers but do not express *Pvalb*^18^. However, so far there has been limited knowledge about the specificity, the degree of overlap between different markers and the total number of distinct GABAergic populations in the MSDB.

Our analysis revealed that MSDB GABAergic neurons can be best described by six subclusters blending into each other. Surprisingly, we found that in the MSDB, several classical GABAergic markers such as *Gad1*, *Gad2,* and *Calb2* are not specific for GABAergic neurons, as high expression profiles of these markers have been found also in the cholinergic and glutamatergic populations. Thus, using Cre dependent lines with these promoters in the MSDB may lead to targeting a broader and less specific population of neurons than anticipated.

One subcluster of the GABA_Lhx6 class, GABA_Lhx6_Pvalb, is characterized by the expression of *Pvalb*. We show that the GABA_Lhx6_Pvalb subcluster is characterized by marker genes whose expression levels are strongly correlated with previously described electrophysiological properties of *Pvalb*^+^ neurons, further validating our transcriptomic analysis. One of these genes is *Hcn1*, the hyperpolarization-activated, cyclic nucleotide-gated (HCN) channel. *Hcn1* is known for its role in the pacemaking activity widely reported in MSDB *Pvalb^+^*neurons, which is crucial for the synchronization of hippocampal theta activity^16,60^.

We further identified the GABA_Lhx6_Npas1 subcluster, which is characterized by the highest expression of somatostatin (*Sst*), a well-characterized marker for GABAergic neurons in the cortex and in the hippocampal formation^103^. While earlier studies reported only scattered *Sst* expression in the MSDB^104^, recent research demonstrated a role of these neurons in appetitive learning^21^ and spatial working memory^105^ highlighting that scattered and sparse expression does not *per se* preclude a behavioral relevance. Notably, these GABA_Lhx6_Npas1 neurons also express the neuronal PAS domain 1 (*Npas1*) gene, which is known to limit interneuron proliferation and is linked to the formation of *Sst* neurons during cortical development^106^. *Npas1^+^* GABAergic neurons have been extensively studied in the nearby ventral pallidum, where they are implicated in anxiety^102^ and sleep regulation^107^. This observation suggests that a deeper experimental investigation of *Npas1^+^/Sst^+^*population could lead to novel insights about the role of the MSDB in homeostatic and motivated behaviors.

As a main marker for the GABA_Lhx6_Nts subcluster we identified neurotensin (*Nts)*. *Nts* has been so far studied in relation to feeding behavior in the hypothalamus and LS^108,109^. Although a few studies have reported the presence of *Nts^+^* neurons in the MSDB^104,110,111^, no functional studies have yet been conducted on this neuronal population. This gap of knowledge could be partially due to the subcluster’s spatial distribution, which is quite lateralized in the mid- and posterior-MSDB coronal slices. Notably, ChAT_Gal neurons express the neurotensin receptor 1 (*Ntsr1*). *In vitro* studies in guinea pigs have shown that neurotensin increases the excitability of MSDB cholinergic neurons, potentially impacting hippocampal theta activity^112^ and hippocampal learning and memory processes^52,112^. Future research on the mechanisms by which the GABA_Lhx6_Nts population interacts with hippocampal rhythmogenesis and plasticity via neurotensin release hold promise to shed new light on non-canonical ways of neuronal communication in the septo-hippocampal system.

Finally, we detected a subcluster expressing high levels of the calcium binding protein calretinin (*Calb2*), GABA_Lhx6_Calb2. Notably, we observed comparable expression levels of *Calb2* in the VGluT2::Septum population. This finding is in line with developmental studies indicating that at E17.5 the VGluT2 MSDB neuron distribution pattern overlaps with *Calb2* but not with *Gad1*^113^.

### Limitations

It is important to point out that transcriptomic studies have intrinsic methodological limitations. First, some cell types may be particularly sensitive to the mechanical procedures required for scRNA-Seq, leading to underrepresentation in the sequencing data as shown by the different proportion of scRNA-Seq and MERFISH cell count for cholinergic and GABAergic neurons. Additionally, MERFISH data acquisition necessitates the *a priori* selection of a panel of genes of interest. When using publicly available datasets, this requirement can constrain the depth of analysis, as it depends heavily on the initially selected genes. For instance, many genes of interest highlighted in our dot plot analysis are absent from the MERFISH dataset, limiting our ability to explore their spatial localization. Lastly, transcriptomic analysis provides only a snapshot of the genes expressed at the time of tissue collection. While these data are valuable for planning future experiments, they need to be further corroborated by protein expression analysis.

Finally, special care should be taken when designing experiments involving transgene expression under specific Cre promoters: gene expression undergoes temporal dynamics over the animal lifetime, and it is essential to reach appropriate levels of the transcript to guarantee a successful and cell-specific neuronal transduction.

### Conclusions and future research

By combining the strengths of both scRNA-seq and MERFISH techniques, we provide the first thorough transcriptomic characterization of MSDB neuron types. Our scRNA-Seq dataset and our comprehensive analysis represent a solid framework to study the MSDB and its diverse cell populations in the future. The discovery that both ChAT and GABAergic neurons are organized in a continuum of subclusters necessitates the development of new approaches, such as intersectional gene targeting and multi-omics assays, to fully understand the functional roles of these neurons at both the molecular and network level. Our finding that the MSDB GABAergic population is ill-represented by widely used GABAergic markers such as *Gad1* and *Gad2* urge future research to carefully select the Cre line’s promoters to target the neuronal population of interest within the medial septal region. Finally, we highlighted how neuromodulators and their receptors have specific expression in neuronal subclusters of the MSDB. The study of galanin and neurotensin effects in the local network and long-range projections towards the hippocampal formation represents a promising avenue to further understand how septal activity is modulated and how, in turn, it influences hippocampal rhythmogenesis and firing patterns. Taken together, our results provide a foundation for developing novel hypotheses to test MSDB cell-type specific contribution in modulating animal behavior. These insights offer promising avenues to understand how MSDB neuron subclusters affect the local and global network in both physiological and pathological conditions.

## AUTHOR CONTRIBUTIONS

Conceptualization: S.P., M.S.C., N.S. and S.R.; Software & Formal analysis: F.K.; Resources: L.W., A.L.L., M.S.C., N.S.; Writing: P.M., F.K. with help from O.B. and L.S.; Visualization: F.K., P.M.; Funding acquisition: S.R.

## SUPPLEMENTAL INFORMATION

Document S1 – Figures S1-S7

## METHODS

### Key resources table

| REAGENT or RESOURCE | SOURCE | IDENTIFIER |
| --- | --- | --- |
| **Deposited data** |  |  |
| scRNA-Seq data | This publication | GEO: GSE163476 |
| Single-cell spatial transcriptomics data | Zhang et al.<sup>40</sup> | N/A |
| Electrophysiological correlates of gene expression | Bomkamp et al.<sup>63</sup> | N/A |
| **Experimental models: Organisms/strains** |  |  |
| C57BL/6 mice | Jackson Laboratory | RRID: IMSR_JAX:000664 |
| **Software and algorithms** |  |  |
| R | R Development Core Team<sup>114</sup> | RRID:SCR_001905 |
| Seurat 5 | Hao et al.<sup>115</sup> | RRID:SCR_007322 |
| Python 3 | van Rossum et al.<sup>116</sup> | RRID:SCR_008394 |
| scanpy | Wolf et al.<sup>117</sup> | RRID:SCR_018139 |
| statsmodels | Seabold et al.<sup>118</sup> | RRID:SCR_016074 |
| PySal-esda | Rey et al.<sup>119</sup> | N/A |
| scMiko | Mikolajewicz et al.<sup>32</sup> | N/A |
| Code | This paper |  |

## EXPERIMENTAL MODEL

### Animals

Experiments used 4 male wildtype C57BL/6 mice between 2-4 months of age, with mice individually housed and maintained on a 12-h light-dark cycle with *ad libitum* access to food and water, with tissue harvested during the light phase. All experimental procedures were approved by the Janelia Research Campus Institutional Animal Care and Use Committee.

## METHOD DETAILS

### scRNA-Seq data acquisition

Single-cell RNA sequencing datasets were generated according to a previously published protocol^28^. In total, four mature C57BL/6 WT male mice were used in to construct datasets (n=2 mice per batch). From each mouse, coronal brain sections were cut, and the medial septum and nucleus of the diagonal band micro-dissected and dissociated. For manual purification of individual cells^120^, cells were aspirated in capillary needles in approximately 0.1-0.5 mL ACSF cocktail and placed into 8-well strips containing 3 µL of cell collection buffer (0.1% Triton X-100, 0.2 U/µL RNAse inhibitor (Lucigen, Middleton, WI), and generally processed according to published methodology^121^.

In brief, each strip of cells was flash frozen on dry ice, then stored at -80°C until cDNA synthesis. Cells were lysed by adding 1 µL lysis mix (50 mM Tris pH 8.0, 5 mM EDTA pH 8.0, 10 mM DTT, 1% Tween-20, 1% Triton X-100, 0.1 g/L Proteinase K (Roche), 2.5 mM dNTPs (Takara), and ERCC Mix 1 (Thermo Fisher) diluted to 1e-6) and 1 µL 10 µM barcoded RT primer (E3V6NEXT primer from ^122^, modified to add a 1 bp spacer before the barcode, extending the barcode length from 6bp to 8bp, and designing the 384 barcodes to tolerate 1 mismatch error correction). The samples were incubated for 5 minutes at 50°C to lyse the cells, followed by 20 minutes at 80°C to inactivate the Proteinase K. Reverse transcription master mix (2 µL 5× buffer (Thermo Fisher Scientific), 2 µL 5M Betaine (Sigma-Aldrich, St. Louis, MO), 0.2 µL 50 µM E5V6NEXT template switch oligo (Integrated DNA Technologies, Coralville, IA), 0.1 µL 200 U/µL Maxima H-RT (Thermo Fisher Scientific), 0.1 µL 40 U/µL RNasin (Lucigen), and 0.6 µL nuclease-free water (Thermo Fisher Scientific) was added to the approximately 5.5 µL lysis reaction and incubated at 42°C for 1.5 hour, followed by 10 minutes at 75°C to inactivate reverse transcriptase. PCR was performed by adding 10 µL 2× HiFi PCR mix (Kapa Biosystems) and 0.5 µl 60 µM SINGV6 primer with the following conditions: 98°C for 3 minutes, 20 cycles of 98°C for 20 seconds, 64°C for 15 seconds, 72°C for 4 minutes, with a final extension step of 5 minutes at 72°C. Samples were pooled across the plate to yield approximately 2 mL pooled PCR reaction. From this, 500 µL was purified with 300 µL Ampure XP beads (0.6× ratio; Beckman Coulter, Brea, CA), washed twice with 75% ethanol, and eluted in 20 µL nuclease-free water. The cDNA concentration was determined using Qubit High-Sensitivity DNA kit (Thermo Fisher Scientific).

Twelve plates were analyzed in total, with six hundred pg cDNA from each plate of cells used in a modified Nextera XT (Illumina, San Diego, CA) library preparation, but using the P5NEXTPT5 primer and extending the tagmentation time to 15 min. The resulting libraries were purified according to the Nextera XT protocol (0.6× ratio) and quantified by qPCR using Kapa Library Quantification (Kapa Biosystems). Plates were pooled on a NextSeq 550 high-output flowcell with 26 bases in read 1, 8 bases for the i7 index, and 50 bases for read 2. Alignment and count-based quantification of single-cell data was performed by removing adapters, tagging transcript reads to barcodes and UMIs, and aligned to the mouse genome via STAR^123^.

### Data preprocessing and batch integration of scRNA-Seq data

For the computational analysis of the scRNA-Seq data the R package Seurat (v.5.0.1)^115^ was used. All functions mentioned hereafter are from the Seurat package with default parameters unless stated otherwise. A Seurat Object for each of the two experimental batches was created separately from the raw count data using the *CreateSeuratObject* function. Only high-quality neurons, which expressed at least 2500 features, at least 10,000 RNAs, and had a count for the neuronal marker *Snap25* of at least 10, were considered. To normalize the data, scale the data, find variable features per cell, and remove confounding sources of variation, the *SCTransform* function was applied to each batch separately. After that, 4000 integration features were determined using the function *FindIntegrationFeatures*, based on which the integration of both batches was conducted using Canonical correlation analysis (CCA)^42^ with the *PrepSCTIntegration*, *FindIntegrationAnchors* and *IntegrateData* functions.

### Dimensionality reduction and (sub-)clustering of scRNA-Seq data

To cluster the data into coarse neuron classes, the dimensionality of the integrated Seurat object was reduced to thirteen dimensions by running principal component analysis (PCA). Thirteen dimensions in principal component (PC) space were selected for clustering based on *ElbowPlot* inspection. The number of dimensions was chosen to be on the higher end because *SCTransform* allows for the identification of more nuanced biological differences and variations across different cell types while maintaining low sensitivity to technical noise and the selected number of dimensions. Then the data was clustered in the dimensionality reduced space with the *FindNeighbors* (dims = 1:13) and *FindClusters* (resolution = 0.1) functions, using the Leiden algorithm^31^. The clustering’s insensitivity to the number of dimensions was demonstrated by reclustering the data using 20 dimensions without change in cell class assignments across all neurons. The resolution for the Leiden clustering of 0.1 was determined using the *multiCluster* function followed by the *multiSpecificity* function of the Seurat-based package *scMiko*^32^. This function clusters the data over a wide range of resolutions and calculates an aggregate score for marker gene specificity for each resolution, based on the co-dependency index (CDI) metric. The optimal resolution was chosen as the one with the highest aggregate score. The identification of glutamatergic, GABAergic and cholinergic clusters was based on the expression profiles of the cholinergic marker gene Chat, the gene for the glutamate transporter VGluT2 (*Slc17a6*) and the gene for the vesicular GABA transporter VGAT (*Slc32a1*). To visualize the clustering results we projected the data points from the reduced feature space on a two dimensional plane using Unifold Manifold Approximation and Projection (UMAP)^123^ with *RunUmap* (dims=1:13).

For subclustering of the GABA_Lhx6 and the ChAT cell class, neurons belonging to the specific class of interest were batch-integrated anew. This new batch integration was performed to disregard effects from non-related neuron classes. For the integration, 2000 integration features were used. Subclustering followed the same procedure as clustering for main cell classes. Both the ChAT class and the GABA_Lhx6 class were subclustered in the first 10 dimensions after PCA using resolutions of 0.4 and 1.2, respectively.

### Differential expression analysis and identification of marker genes

To identify the top differentially expressed genes (DEGs) for all four coarse cell classes, the Seurat object’s SCT assay was used. Initially, the *PrepSCTFindMarkers* function was run, followed by the Wilcoxon rank sum test for DEG identification using *FindMarkers* (*min.diff.pct* = 0.3, *logfc.threshold* = 1.5, *only.pos* = TRUE). The test compared the expression of each gene in a certain class to its expression in all other classes combined. This was also done for DEG identification of ChAT and GABA_Lhx6 subclusters, only that here the test compared gene expression in the respective subcluster with its expression in all other subclusters of the respective class combined. The min-max scaled log-expression of the 10 DEGs with the lowest FDR-corrected p-values (using Bonferroni correction) per cluster was visualized on heatmaps using the *DoHeatmap* function. To illustrate the continuous character of the ChAT subclusters, the expression value of selected DEGs was standardized and sorted according to their rank on the diffusion pseudotime axis before being plotted on a heatmap.

The *R* package *EnhancedVolcano*^124^ was employed to display DEG expression between the ChAT_Amigo2 and ChAT_Gal subclusters in a Volcano plot. Exclusive marker genes for subclusters were identified by calculating CDI and CDI based p-values for each gene using the *findCDIMarkers* function of the *scMiko* package. The function was run on the Seurat object’s SCT assay after running *PrepSCTFindMarkers*. For the suffixes of the subclusters we used genes which were previously associated with neuromodulatory function (*Adra1a*^125^ for GABA_Lhx6_Adra1a), or common Cre-lines (*Amigo2* for ChAT_Amigo2 - Jackson Strain #:030215; *Gal* for ChAT_Gal; *Nts*^112^ for GABA_Lhx6_Nts - Jackson Strain #:017525; *Calb2*^18^ for GABA_Lhx6_Calb2 - Jackson Strain #:010774; *Pvalb*^64^ for GABA_Lhx6_Pvalb, Jackson Strain #:017320). Additionally, the markers used for the suffices were checked to all have a normalized CDI of at least 0.2 and an adjusted p-value below 1e-6. If no previous association was apparent with any of the top marker genes, we named the subclusters by using genes with the lowest p-value that had a normalized CDI of at least 0.2 as suffixes (*Tcerg1l* for GABA_Lhx6_Tcerg1l, *Npas1* for GABA_Lhx6_Npas1.)

### Subcluster discreteness analysis

To evaluate the discreteness of GABA_Lhx6 and ChAT subclusters, a method introduced by Stanley and colleagues^53^ was employed to examine gaps in diffusion pseudotime and relative cluster connectivity. Diffusion pseudotime, which places cells on an axis reflecting the highest variance between two clusters^54^, and relative cluster connectivity, which measures a cell’s similarity to cells within the same cluster compared to those in different clusters, were both calculated. Discrete clusters exhibit a larger gap in diffusion pseudotime compared to continuous ones. This gap was quantified using the gap c statistic. For the analysis of ChAT and GABA_Lhx6 subclusters, diffusion pseudotime, relative cluster connectivity, and the gap c statistic were calculated using customized code based on code from Stanley et al.^53^ which relies on the *scanpy* package^117^.

As an additional assessment of ChAT subcluster discreteness, the relative gene score of each cell was plotted as a function of pseudotime rank. To calculate the relative gene score for each ChAT cell, the top 20 positive DEGs for both subclusters were calculated using the Wilcoxon rank sum test. It was ensured that all of them had an FDR corrected p-value of below 10^-6^. To calculate a cumulative score, the log-transformed z-scored gene expression of positive DEGs for the ChAT_Amigo2 subcluster was added, and the log-transformed z-scored gene expression of positive DEGs for the ChAT_Gal subcluster was subtracted for each cell individually.

### Correlation with electrophysiological properties

To identify how differentially expressed genes (DEGs) of the GABA_Lhx6_Pvalb subcluster correlate with electrophysiological properties, positive DEGs for GABA_Lhx6_Pvalb neurons compared to all other neurons of the GABA_Lhx6 class were calculated using Wilcoxon rank sum test with *FindMarkers* (*min.pct* = 0.2 and *logfc.threshold* = 0.5). They were further subsetted for genes with a maximum FDR corrected p-value of 1e-4. The database from Bomkamp and colleagues^63^ was then utilized to identify DEGs which had significant correlations (either positive or negative) with electrophysiological properties in inhibitory neurons. A significance threshold of *FDR_gene_inh_only* < 0.01 was applied. Next, a heatmap was constructed showing the log2 fold-change expression of significantly upregulated DEGs of the GABA_Lhx6_Pvalb cluster which also had a significant correlation with electrophysiological properties. Electrophysiological properties were included in the heatmap only if they had either four times as many positively correlated genes as negatively correlated genes, or four times as many negatively correlated genes as positively correlated genes. For GABA_Lhx6 neurons not belonging to the GABA_Lhx6_Pvalb subcluster, the same analysis was performed with negative DEGs of the GABA_Lhx6_Pvalb subcluster (i.e., genes significantly downregulated in GABA_Lhx6_Pvalb).

### MERFISH data integration and analysis

For the spatial transcriptomics analyses, two coronal MERFISH datasets, Zhuang-ABCA-1 and Zhuang-ABCA-2^40^, from the Allen Brain Cell Atlas (ABCA) were utilized. Both datasets included 1122 genes. The segmentation of mRNA spots into single cells was already given by the Zhuang-ABCA datasets and adopted. The segmentation of the MSDB was based on alignment with the Allen Mouse Brain Common Coordinate Framework version 3 (CCF)^41^, which was also provided with the Zhuang-ABCA datasets. The MSDB segmentation was further refined and cropped to include only the left hemisphere using a custom-made tool. This refinement was done to avoid any bias in cell quantification due to the incomplete representation of the right hemisphere in both Zhuang datasets. Both Zhuang datasets were further subset to include only neurons. For each dataset a separate Seurat object was created and the *SCTransform* function was applied to both Seurat objects separately. Subsequently, CCA-based integration of both objects was performed using the *FindIntegrationAnchors* and *IntegrateData* functions.

### MERFISH cell-type prediction and validation

To predict the class identity of MERFISH neurons based on the scRNA-Seq analysis, CCA-based prediction scores were calculated for all MERFISH neurons using Seurat’s *FindTransferAnchors* (dims = 1:13) and *TransferData* (dims = 1:13) functions. The integrated MERFISH dataset served as the query, while the cell class identities of the integrated scRNA-Seq neurons served as reference. Only neurons with a maximum prediction score above 0.9 were assigned class labels. To investigate the spatial distribution of cell types within the MSDB and in its periphery, we conducted this analysis on cells within a selection window that encompassed all coronal slices containing parts of the MSDB. The selection window extended from CCFz = 4 to CCFz = 6, and from CCFy = 3 to CCFy = 7 on each slice, using the coronal slice coordinates in the Common Coordinate Framework (CCF). For all the analyses mentioned hereafter, the term *selection window* refers to the window described here.

To validate the predicted class identity of the MERFISH neurons, marker genes for the predicted MERFISH cell classes were identified with *scMiko*’s *findCDIMarkers* function. Gene expression profiles displaying the min-max scaled log-expression of the top 10 marker genes which had the lowest p-value, a normalized CDI score of at least 0.2 and which were present in both scRNA-Seq and MERFISH datasets were confirmed for similarity between scRNA-Seq and MERFISH datasets.

To ensure that no major cell class was overlooked in the scRNA-Seq dataset, an integrated Seurat object of MSDB^+^ neurons from both MERFISH datasets and both scRNA-Seq batches was created using the *FindIntegrationAnchors* (dims = 1:13) and *IntegrateData* (dims = 1:13) functions. A UMAP was then computed in the integrated dataset’s feature space (dims = 1:13), referred to as joint feature space (jFS). The similar distribution pattern of both MERFISH and scRNA-Seq neurons on the UMAP indicated that no major neuron class was missed in the scRNA-Seq dataset.

For the prediction and integration of MERFISH ChAT subclusters, the same analysis as for the main neuron classes was employed with a few differences: only MERFISH neurons previously predicted as ChAT neurons were used for prediction and integration; both prediction and integration were conducted in the first 10, instead of the first 13 dimensions of PC space; and the threshold for prediction scores was lowered to 0.5. This adjustment ensured that cells lying in the middle of the continuum between both ChAT subclusters were not disregarded. The analysis was repeated, including MERFISH ChAT neurons from both MSDB and Striatum by selecting cells from the selection window. After subclustering the integrated dataset of MSDB and Striatal ChAT MERFISH neurons (following the previously described subclustering workflow - here with dims = 1:13 and a Leiden resolution of 0.05), it was found that Striatal ChAT neurons formed a distinct cluster compared to MSDB^+^ ChAT neurons. Therefore, subcluster identities from scRNA-Seq ChAT subclusters were only predicted for MERFISH ChAT neurons belonging to the cluster populated with MSDB^+^ cells. Neurons from the other cluster were assigned the label ChAT::Striatum.

The subcluster identities of MSDB^+^ MERFISH neurons of the GABA_Lhx6 class were determined using the same methods as for the ChAT class. Prediction scores for MERFISH GABA_Lhx6 neurons, as well as the subcluster identities of GABA_Lhx6 scRNA-Seq neurons, were visualized on a UMAP in jFS. Since some regions of the jFS were only sparsely populated by scRNA-Seq neurons, but densely by MERFISH neurons, it was checked if the MERFISH neurons in this region constitute separate subclusters that were not represented in the scRNA-Seq dataset. Therefore, the GABA_Lhx6 neurons from both MERFISH datasets were additionally integrated without the scRNA-Seq neurons and subclustered using the same functions for integration and subclustering as for the GABA_Lhx6 scRNA-Seq data; here with a Leiden resolution of 0.3. The MERFISH subcluster assignments were also visualized in the jFS. The areas of the jFS which were densely populated by MERFISH neurons and sparsely by scRNA-Seq cells did not constitute separate MERFISH clusters, indicating that the scRNA-Seq data covered the spectrum of GABA_Lhx6 subclusters well.

To further validate the prediction for the MERFISH GABA_Lhx6 subclusters, the top marker genes for the predicted subclusters of MERFISH GABA_Lhx6 neurons were identified with *scMiko*’s *findCDIMarkers* function. Heatmaps displaying the min-max scaled expression of the top 10 marker genes which had the lowest p-value, a normalized CDI score of at least 0.2 and which were present in both scRNA-Seq and MERFISH datasets were plotted for both MERFISH and scRNA-Seq neurons and confirmed for similarity. For the spatial analysis (described later in the Methods section), prediction and integration were additionally conducted for neurons in the selection window, also including neurons outside of the MSDB borders. For this all functions and parameters were the same as for the prediction and integration of MSDB^+^ neurons only.

### Clustering analysis of VGluT2 cells in MERFISH datasets

Neurons from inside the selection window were chosen from both MERFISH datasets and further subsetted for VGluT2^+^ neurons only. The selected neurons of both MERFISH datasets were integrated and subclustered as described for the GABA_Lhx6 MERFISH neurons, only that here the first 5 dimensions in PC space and a Leiden resolution of 0.05 was chosen after inspection of the *ElbowPlot*. The relatively low number of PCs and the relatively low resolution focuses the analysis on the most significant sources of variation, minimizing the effects of a potential bias that could have been introduced by the pre-selected and limited MERFISH gene set.

Since the cell number of scRNA-Seq neurons was relatively small and thus not optimal for CCA-based integration with the MERFISH dataset, CDI-based markers were calculated for the glutamatergic MERFISH clusters using *scMiko*’s *findCDIMarkers* function on the integrated dataset. The min-max scaled log-transformed expression of the marker genes with the lowest p-values, which had a normalized CDI score of at least 0.2, were plotted in a heatmap. Additionally, the min-max scaled log-expression of the same genes was plotted for all scRNA-Seq neurons of the VGluT2 class, showing that the expression profile of the scRNA-Seq VGluT2 neurons corresponds with the expression profile of the VGluT2::Septum class.

### Spatial analysis

To show the spatial distribution of MERFISH cell classes and subclusters, cells inside of the selection window were plotted in CCF-coordinates color coded by the specific classes or subclusters of interest. Significant spatial trends were investigated among MSDB^+^ neurons inside of each MERFISH slice, as well as along an anterior-posterior axis across slices.

Significant spatial trends of MSDB^+^ neurons across the anterior-posterior axis were determined by using a Monte-Carlo approach. Class or subcluster labels of MSDB^+^ neurons across all MSDB^+^ slices were shuffled 1,000 times. A neuron class or subcluster was determined to be significantly enriched in a slice along the anterior-posterior axis if it was abundant more often than in 95 percent of the shuffles (p < 0.05). For anterior-posterior trends of neuron classes the shuffling was performed across all neurons; for Chat subclusters and GABA_Lhx6 subclusters, shuffling was performed among ChAT and GABA_Lhx6 neurons only.

Spatial trends of MSDB^+^ neurons along the mediolateral and dorsoventral axes were investigated for each individual slice separately. First, a k-nearest neighbors’ graph was constructed using euclidean distance. To account for the differences in cell densities, a k of 15 was used for main cell classes, a k of 10 was used for GABAergic subclusters and a k of 5 was used for ChAT subclusters. Each edge of the graph that connected two cells of the respective class or subcluster of interest was given a weight of 1; all other edges were given a weight of 0. The graph was then used to calculate global Moran’s I^47^ on it, which is a metric that quantifies the spatial clustering of the data. Moran’s I was calculated using the *moran.Moran* function of the Python package *PySal-esda*^119^. To determine if the spatial clustering observed on the respective slice was significantly above chance, a Monte-Carlo approach was used. Class/subcluster labels of the neurons of interest were shuffled 1,000 times and global Moran’s I was calculated for each of the shuffles. Spatial clustering was determined to be significant if the rank of the actual global Moran’s I was above the 95th percentile (p < 0.05) of all shuffles. For the main neuron classes, all MSDB^+^ neurons of the respective MERISH slice were used for shuffling; for ChAT subclusters, only ChAT neurons were used, and for the GABA_Lhx6 subclusters, only GABA_Lhx6 neurons were used. The procedure was repeated for each class/subcluster separately.

For neuron classes/subclusters that showed significant spatial clustering inside MSDB on individual slices, a two-dimensional maximum-likelihood kernel density estimation (ml-KDE) was calculated using the *KDEMultivariate* function of the Python package *statsmodels*^118^. The two-dimensional full width at half maximum (FWHM) of the ml-KDE was plotted on top of the respective spatial plot.

## ACKNOWLEDGMENTS

We thank the quantitative genomics unit at HHMI’s Janelia Research Campus for the technical support, and the entire S.R. laboratory for insightful discussions. The authors received funding from the German Research Foundation (DFG) CRC 1436 - project 425899996 (A03 - to S.R.), CRC 1089 (to S.R.), RTG 2413 SynAGE project 362321501 (to S.R.), Leibniz SAW Learning Resilience (to S.R.), ERC Consolidator grant Sub-D-Code (to S.R.) and the BMBF DZPG (01EE2305E, to S.R.). The collaboration between S.P., M.S.C., S.R. and N.S. was supported by the HHMI Janelia Visiting Scientist Program.

