## Supplementary figures for "Neuronal heterogeneity in the medial septum and diagonal band of Broca: classes and continua"

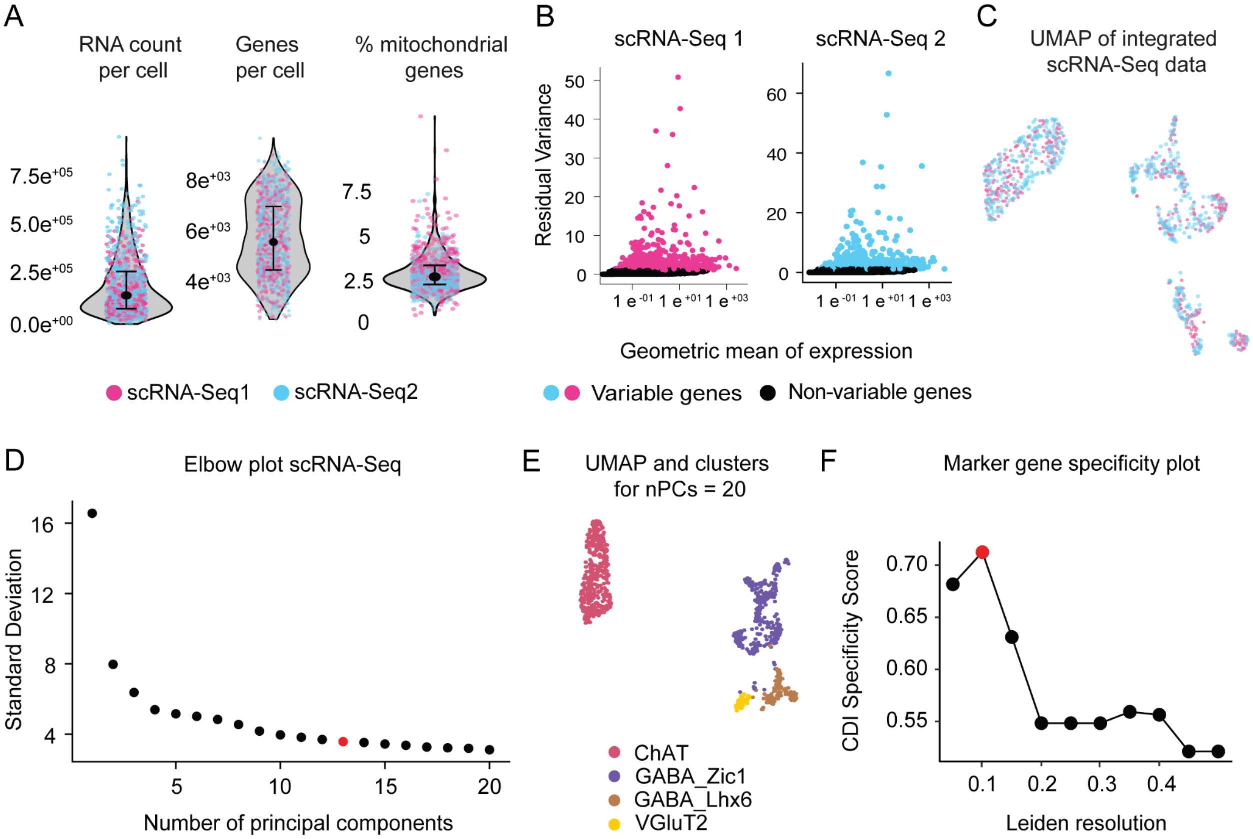


**Figure S1 (related to Figure 1). Quality control for scRNA-Seq data and clustering validation.**
A) Violin plots with RNA count per cell (median = 154711, IQR = 89072-273657), genes (features) per cell (median = 5563, IQR = 4477 -6947) and percentage of mitochondrial genes per cell of MSDB scRNA-Seq neurons (median = 2.7, IQR = 2.26-3.35) color coded by batch identity (scRNA-Seq1 in magenta, scRNA-Seq2 in cyan, 2 mice per batch) after the selection of high quality cells (see Methods). Median and IQR are shown in black in the violin plots. B) Feature variability plots for scRNA-Seq1 and scRNA-Seq2 showing the residual variance of each gene as a function of its mean expression. (Number of genes in batch 1: 16209, number of genes in batch 2: 16768, number of variable genes per batch: 3000; non-variable genes in black). C) UMAP of integrated scRNA-Seq dataset. Batch identity of each cell is color coded (scRNA-Seq1 in magenta, scRNA-Seq2 in cyan). D) Elbow plot showing standard deviation as a function of the number of principal components. 13 PCs were used for further clustering (shown in red). E) UMAP of MSDB clusters calculated with 20 PCs instead of 13. The grouping of the cells is identical to the one with 13 PCs only, showing robustness of the method towards the number of PCs. F) CDI specificity score as a function of Leiden resolution. The highest specificity is reached at a resolution of 0.1 (indicated in red) which was used for Leiden Clustering of the main cell classes.

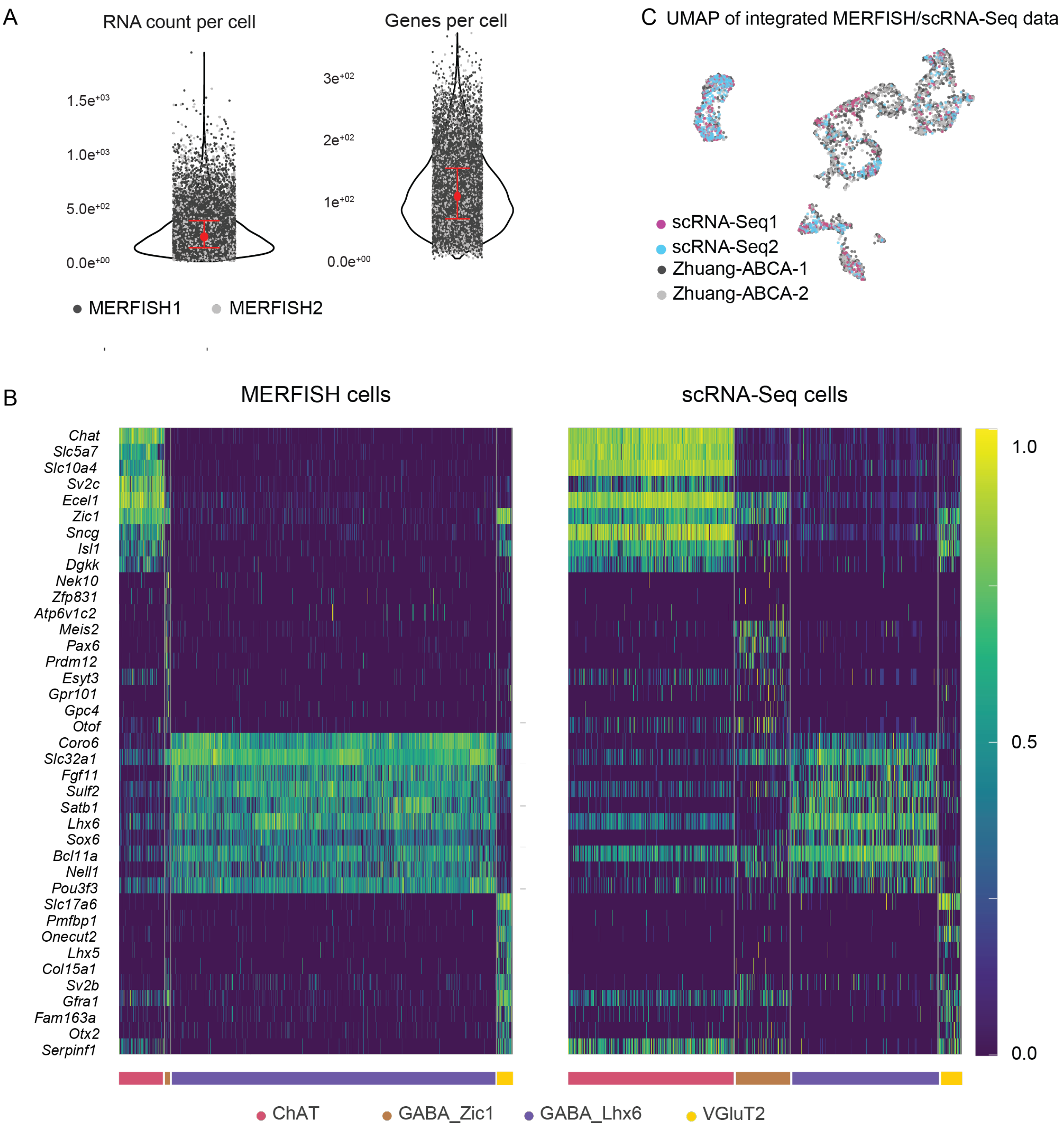


**Figure S2 (related to Figure 2). Validation of scRNA-Seq integration with MERFISH ABCA dataset.**
A) Number of RNAs per cell (median: 240, IQR: 136-395) and number of individual genes per cell (median: 105, IQR: 69-150) for both MERFISH datasets (Zhuang-ABCA-1: MERFISH1 and Zhuang-ABCA-2: MERFISH2). B) *Left:* Heatmap of the log-transformed scaled expression of top marker genes for each predicted class of MERFISH neurons; calculated using co-dependency index-based marker gene identification (see Methods). *Right*: Expression of these genes in scRNA-Seq neurons. C) UMAP in jFS of MERFISH and scRNA-Seq neurons after integration color coded by batch identity (scRNA-Seq1 in magenta, scRNA-Seq2 in cyan, Zhuang-ABCA-1 in gray, Zhuang-ABCA-2 in black).

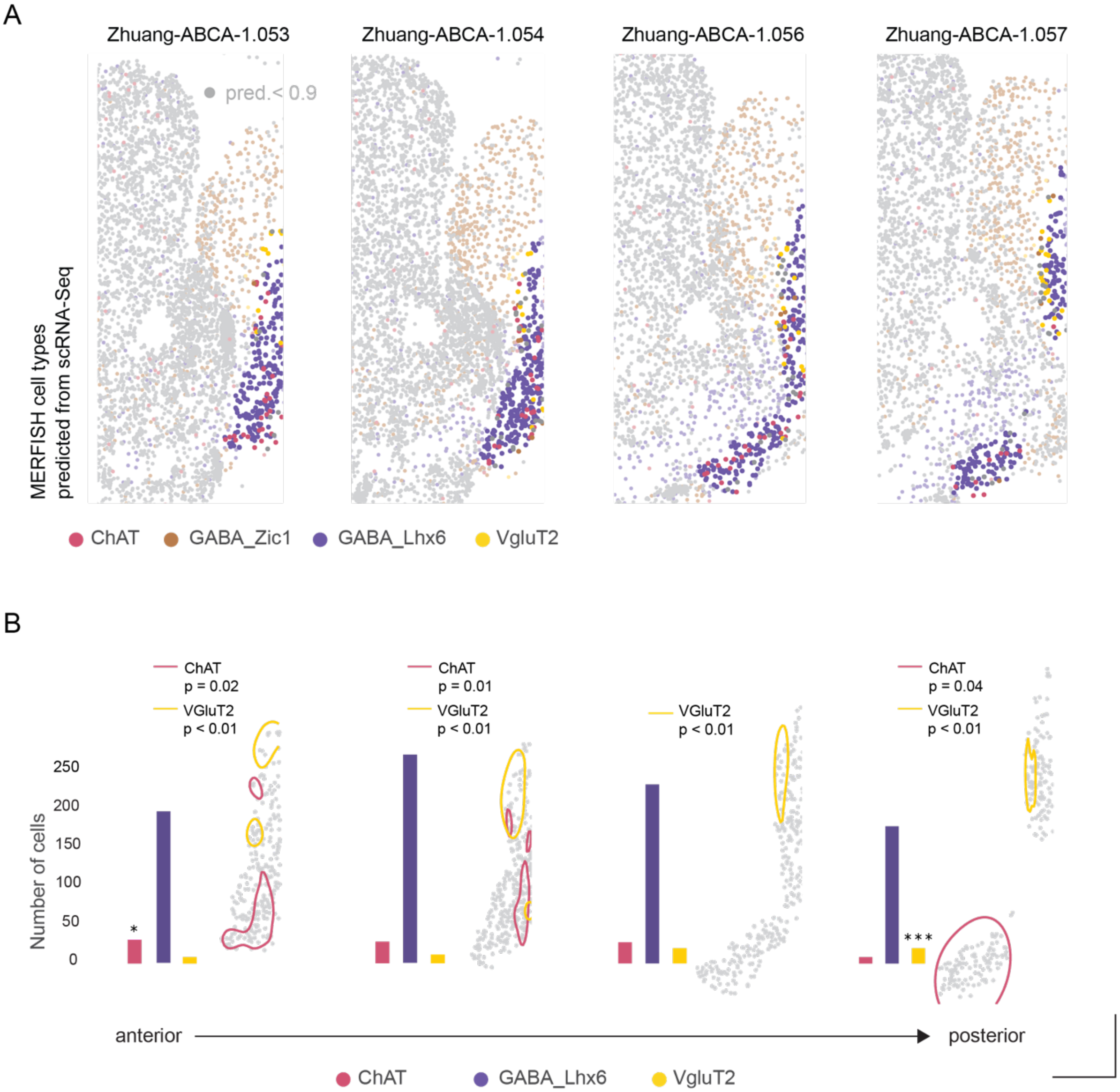


**Figure S3 (related to Figure 2). Spatial distribution of MSDB main cell types in the Zhuang1 dataset.**
A) MERFISH coronal slices of Zhuang-ABCA-1 dataset (slices 053, 054, 056, and 057) with cells color-coded based on the predicted identity from the scRNA-Seq dataset (ChAT in pink, GABA_Zic1 in brown, GABA_Lhx6 in purple, VGluT2 in yellow, MSDB selection in bold colors, outside selection in faint colors). Scale bars correspond to 5 CCF units for both axes. B) Bar plots represent the number of neurons per cluster for each coronal slice. Stars represent significant enrichment of the respective cluster on the respective slice following Monte-Carlo based permutation testing (* $p<0.05$, ** $p<0.01$, *** $p< 0.001$). FWHM of the ml-KDE estimate represent the location of cell types that are significantly spatially localized inside the MSDB in the respective slice. P-values correspond to the permutation test with 1,000 shuffles (see Methods). Scale bars correspond to 5 CCF units for both axes.

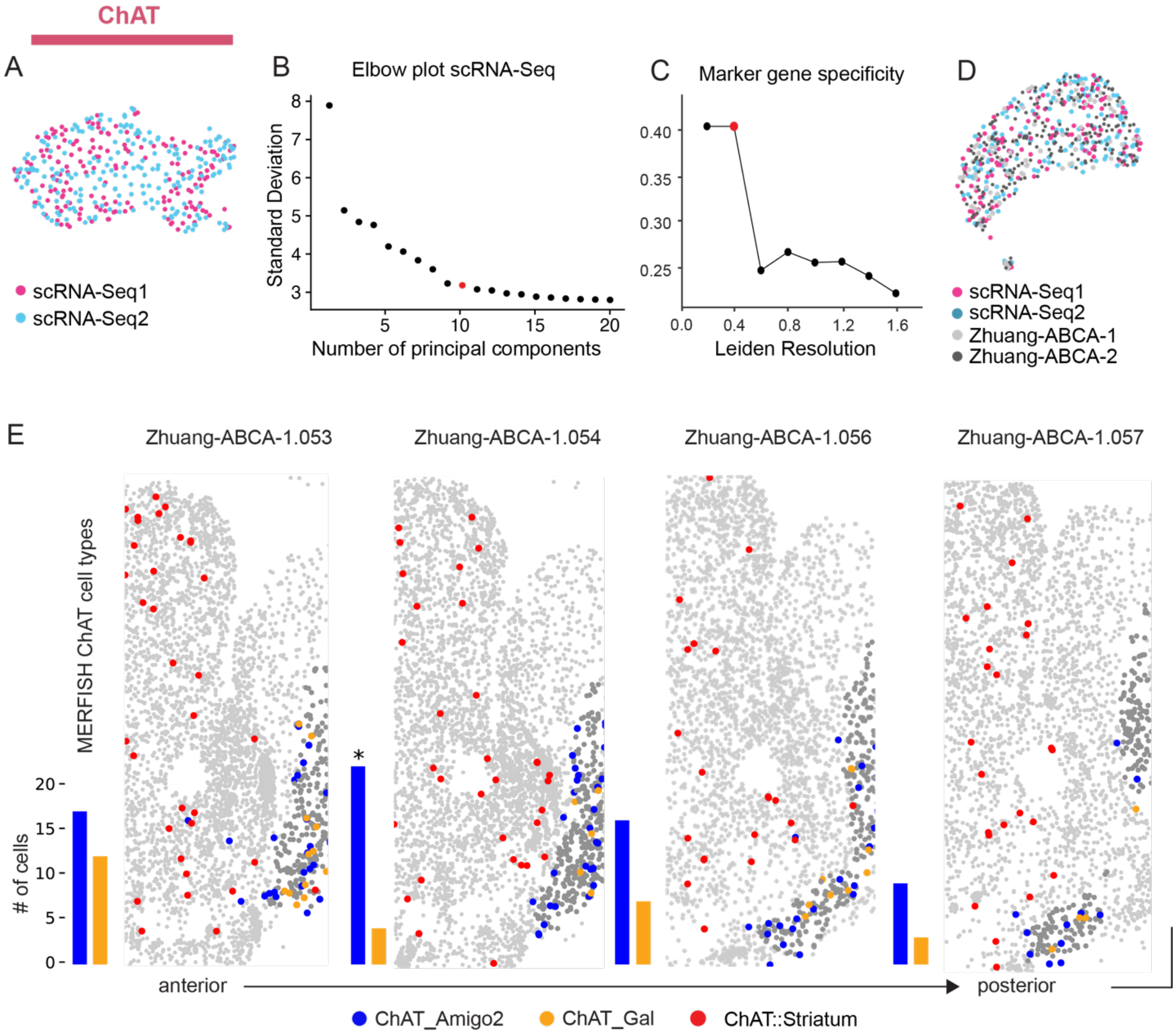


**Figure S4 (related to Figure 3 and 4). Clustering validation of ChAT MSDB neurons and spatial distribution in the Zhuang-ABCA-1 dataset.**

A) UMAP of scRNA-Seq cholinergic neurons color coded by batch identity after batch integration (scRNA-Seq1 in magenta, scRNA-Seq2 in cyan). B) Elbow plot showing standard deviation as a function of the number of principal components for ChAT scRNA-Seq neurons. 10 PCs were used for further clustering (shown in red). C) CDI Specificity Score as a function of Leiden resolution. The highest specificity is reached at a resolution of 0.4 (indicated in red) which was used for Leiden Clustering of the ChAT class. D) UMAP in jFS showing the integration of the two scRNA-Seq batches and the two MERFISH datasets (scRNA-Seq1 in magenta, scRNA-Seq2 in cyan, Zhuang-ABCA-1 in gray, Zhuang-ABCA-2 in black). E) Spatial distribution of the cholinergic clusters along the anterior-posterior axis. (ChAT_Amigo2 in blue, ChAT_Gal in orange, ChAT::Striatum in red). Bar plots represent the abundance of cholinergic neurons from the two MSDB clusters in each slice. Star represents significant enrichment of ChAT_Amigo2 neurons in the Zhuang-ABCA-1.054 slice.

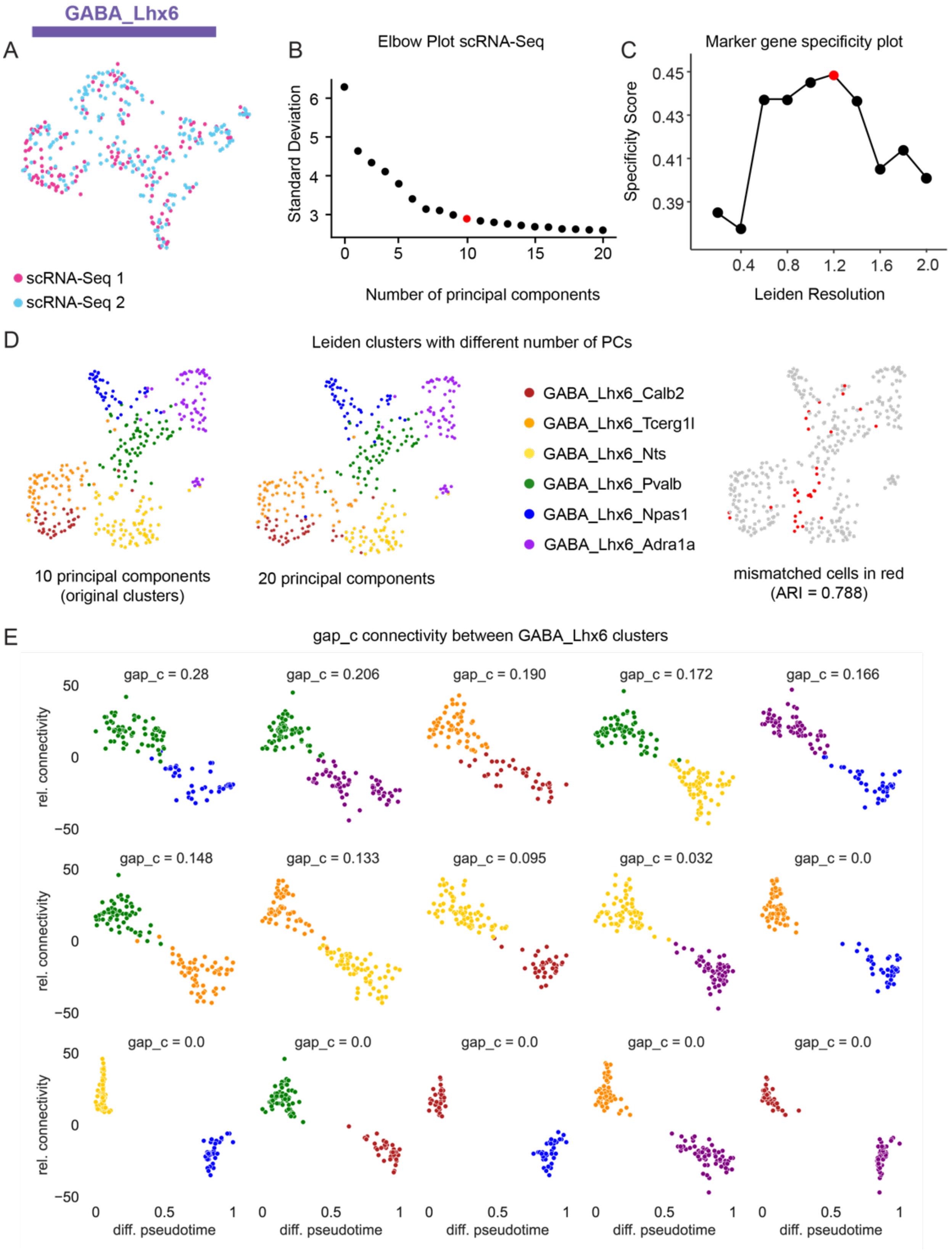


**Figure S5 (related to Figure 5). Validation of MSDB GABA neuron clusters and continuum.**

**Figure S5 (related to Figure 5). Validation of MSDB GABA neuron clusters and continuum (from previous page).** A) UMAP of scRNA-Seq GABAergic neurons color coded by batch identity (scRNA-Seq1 in magenta, scRNA-Seq2 in cyan). B) Elbow plot showing standard deviation as a function of the number of principal components for GABA_Lhx6 scRNA-Seq neurons. 10 PCs were used for further clustering (shown in red). C) CDI Specificity Score as a function of Leiden resolution. The highest specificity is reached at a resolution of 1.2 (indicated in red) which was used for Leiden Clustering of the GABA_Lhx6 class. D) Comparison of GABA_Lhx6 clusters calculated with 10 PCs (right panel) or 20 PCs (middle panel) on a UMAP that was calculated in 20 dimensions. Only few cells are mismatched between the two UMAPs showing robustness of the method towards the number of PCs (adjusted rand index (ARI) = 0.788). E) Relative cluster connectivity of neurons as a function of diffusion pseudotime along the axis of highest variance between the two subclusters sorted by gap c value. The color code represents subcluster identity.

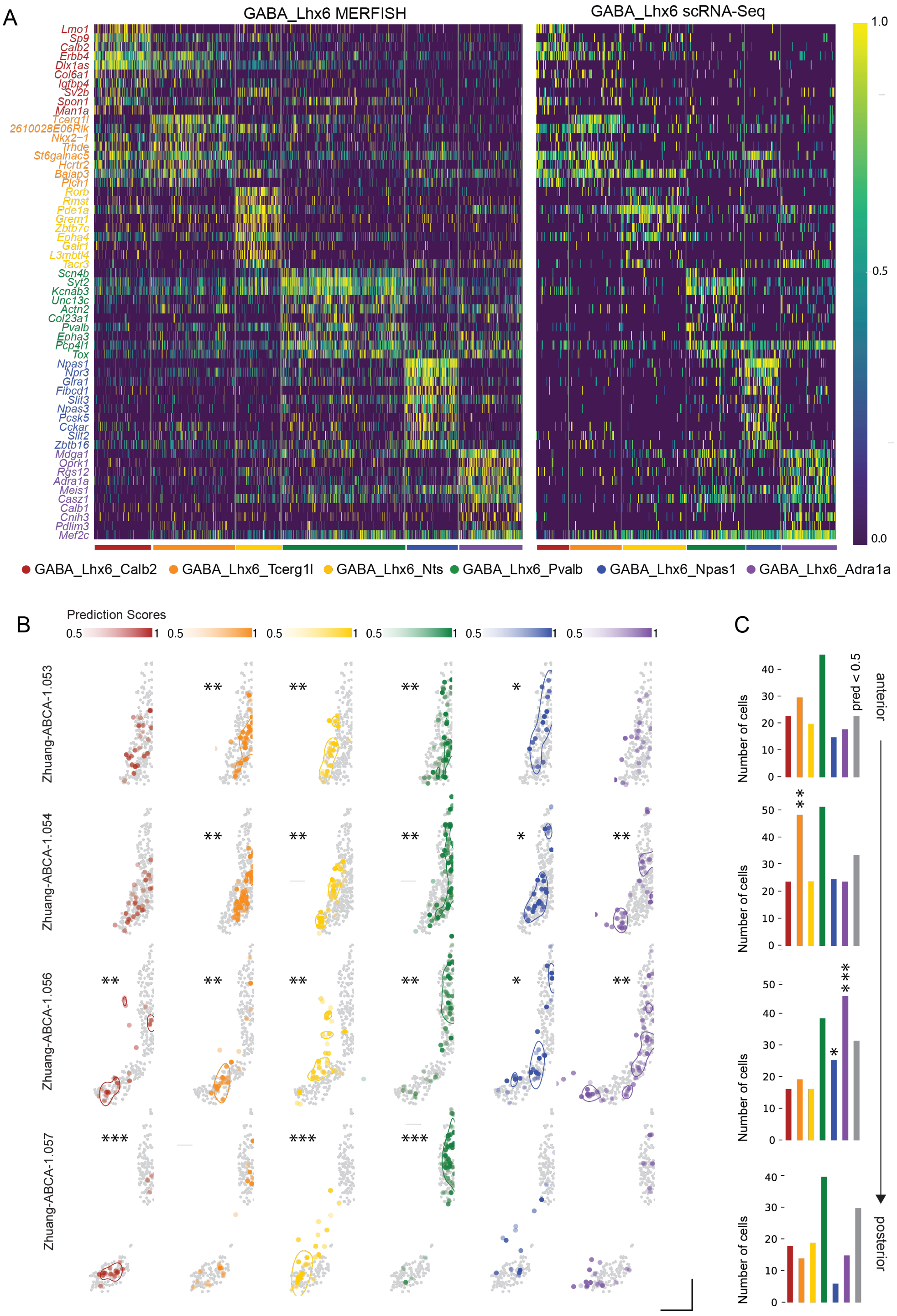


**Figure S6 (related to figure 6). DEG heatmap and spatial distribution of GABA_Lhx6 MERFISH neurons in the Zhuang-ABCA-1 dataset.**

**Figure S6 (related to figure 6). DEG heatmap and spatial distribution of GABA_Lhx6 MERFISH neurons in the Zhuang-ABCA-1 dataset (from previous page).**

A) Heatmap of the log-transformed scaled expression of top marker genes for MERFISH neurons for different GABA_Lhx6 subclusters in both MERFISH neurons and scRNA-Seq neurons. Calculated using co-dependency index-based marker gene identification (see Methods). B) Spatial expression patterns of predicted MERFISH subclusters color-coded by prediction score in the Zhuang-ABCA-1 dataset. Significant spatial clustering inside of the MSDB is represented by FWHM of the respective kernel densities (* $p<0.05$, ** $p<0.01$, *** $p< 0.001$). Scale bars correspond to 5 CCF units for both axes. C) Number of neurons per cluster for each coronal slice. Stars represent significant enrichment of the respective cluster on the respective slice (* $p<0.05$, ** $p<0.01$, *** $p< 0.001$).

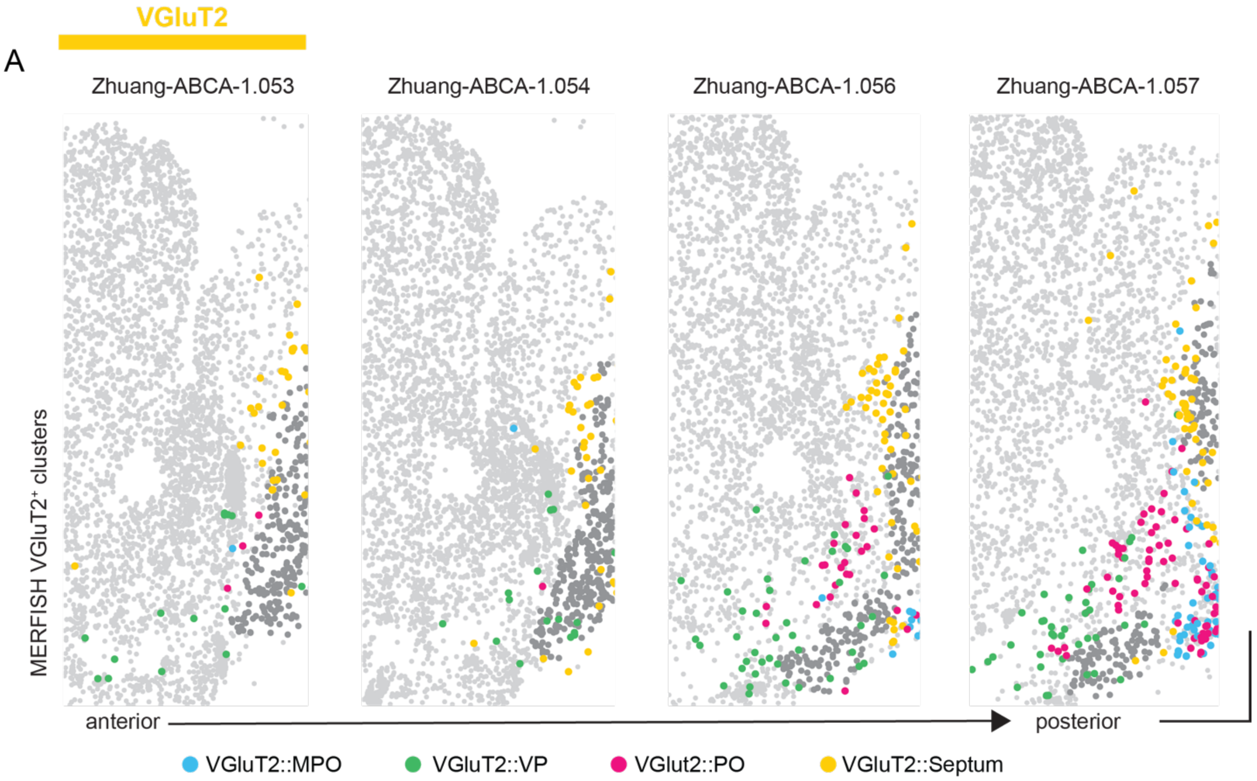


**Figure S7 (related to figure 7). Spatial distribution of VGluT2 MERFISH neurons in the Zhuang-ABCA-1 dataset.** A) Spatial distribution of VGluT2 neurons in the selection window along the anterior-posterior axis of the Zhuang-ABCA-1 dataset. Scale bars correspond to 5 CCF units for both axes.
